# Serosurveillance Studies of Peste des Petits Ruminants (PPR) Virus in Sheep and Goats in South Asia: A Systematic Review and Meta-analysis

**DOI:** 10.1101/2025.06.26.661882

**Authors:** Md Jisan Ahmed, Md Imran Hossain, Ritu Chalise, Md Arifur Rahman, Prajwal Bhandari, Tahmina Sikder, Amina Khatun, Delower Hossain

## Abstract

**Background:** Peste des Petits Ruminants (PPR), also known as the goat plague, is one of the WOAH-listed A, highly contagious and economically important viral transboundary animal diseases affecting small ruminants, and having a significant impact on the global livestock industry and international animal traffic.

**Objective:** The present study aimed to use a systematic approach to assess the pooled seroprevalence of PPRV in sheep and goats in South Asia, through a systematic review and meta-analysis of published data.

**Methods:** A thorough search on various databases was performed to identify published research articles published between January 2000 and June 2025 reporting the seroprevalence of PPRV in small ruminants in South Asia. The articles were chosen on the basis of specific inclusion and exclusion criteria. Since the heterogeneity among the studies was significant, the pooled seroprevalence was estimated via a random effects meta-analysis model, using Stata (v19) and R software (v4.5.0).

**Results:** In sheep and goats, the estimated pooled seroprevalence of PPR was 42.4% (95% CI: 35.0–49.9), whereas it was 41.8% (95% CI: 33.7–50.1) in goats and 44.5% (95% CI: 37.0– 52.0) in sheep. Subgroup analysis revealed that the pooled seroprevalence of PPRV in sheep and goat by country and vaccination status was greater in Nepal (57.0%, 95% CI: 8.4-97.7) and in vaccinated animals (57.5%, 95% CI: 47.9-66.9).

**Conclusion:** This study highlights the need for coordinated actions, including vaccination, surveillance, and strict biosecurity, to control and eradicate the disease effectively. Moreover, authorities should adopt evidence-based strategies to support the global goal of eradicating PPR by 2030, as recommended by the WOAH.

## 1. Introduction

*Peste des petits ruminants* (PPR) is a highly contagious and economically important viral transboundary animal diseases (TADs) that affect small ruminants predominantly in sheep and goats, and pose a significant threat to the health and productivity of these animals. The disease is also known by several other names, such as ‘Goat Plague’, ‘Ovine Rinderpest’, ‘Kata’, and ‘Stomatitis–Pneumonitis Syndrome’ [1]. The World Organization for Animal Health (WOAH, former OIE) has categorized PPR as list-A diseases because of its highly infectious characteristics [2]. PPR is caused by the *peste des petits ruminants virus* (PPRV), which is classified under the *Morbillivirus* genus and belongs to the *Paramyxoviridae* virus family [3]. PPR was first identified in the 1940s in Ivory Coast, West Africa, and has since spread worldwide, now affecting more than 70 countries (according to WOAH), with more than 20 considered at risk on the basis of epidemiological data [4,5]. Over 90% of the world’s small ruminants, including sheep and goats, are in developing countries, where they support food security, trade, and the livelihoods of small-scale farmers [6]. However, sustainable small-ruminant farming, especially sheep and goat farming, is always challenging in developing regions, particularly in Asia, due to the disease, known as PPR. In South Asia, PPR was first detected in southern India in 1987, in Bangladesh in 1993, in Pakistan in 1994, and in Nepal in 1995 [7,8]. PPR causes huge economic losses estimated at approximately USD 3,012.6 million annually in South Asian countries [7].

The variation in PPR sero-prevalence may be attributed to several factors, including the occurrence of PPRV outbreaks in specific regions, differences in diagnostic methods, the source and selection of samples, sampling techniques, the time period, and the particular animal species examined [9]. In many studies, the sero-prevalence of PPRV has been reported to range from 30.91% to 54.2% in India [10], whereas in Bangladesh, it typically ranges between 20% and 30% [8]. Despite the variations in epidemiological, diagnostic, and vaccine production capacities across South Asian countries, Bangladesh, India, Nepal, and Pakistan have demonstrated a strong commitment to implement national action plans aligned with the global PPR eradication campaign [11].

PPRV transmission occurs primarily through aerosols and direct contact between infected and susceptible animals [12]. The incubation period of PPR typically ranges from 4-6 days but can last up to 14 days. Clinical signs include fever, oculo-nasal discharge, eroded lesions of the oral mucosa, diarrhea, and pneumonia [13]. PPRV usually causes high morbidity and mortality, sometimes reaching up to 100% among susceptible non-vaccinated hosts [14]. Disease severity is influenced by various factors, including host species, age, sex, breed, immune status, and previous exposure [15]. Small ruminants, especially sheep and goats in endemic regions, often develop lifelong immunity after infection, whereas naïve populations contribute to the continued circulation of the virus [16]. PPR diagnosis in sheep and goats ranges from identifying clinical symptoms to virus isolation, whereas virus isolation is considered the gold standard [17]. Effective PPR control depends largely on the early detection of infected animals, while serological tests play key roles in identifying subclinical cases [18]. PPRV is antigenically similar to the viruses that cause rinderpest in cattle, measles in humans, and distemper in dogs, however, it can be serologically identified and differentiated using commercially available ELISA kits [19]. Serological surveys can identify immunological gaps by highlighting regions with low immunity levels, enabling targeted vaccination efforts to address these vulnerable populations effectively [20]. As a result, mass vaccination has become a key strategy for controlling PPR and preventing further spread of the disease in the locality. In endemic countries, live attenuated PPRV vaccines are commonly used and offer long-term immunity in sheep and goats [21]; these vaccines generate maternal antibodies in colostrum that are detectable in offspring until three to six months of age and pose no risk to pregnant sheep or goats at any gestational phase [22]. However, ineffective vaccine allocation could contribute to supply shortages. Inadequate data gathering and analytical capabilities hinder evidence-based vaccine distribution and strategic deployment based on epidemiological needs [23]. Therefore, the current study was designed to conduct a systematic review and meta-analysis of the reported serological studies on PPR in South Asian countries. To the best of our knowledge, no reviews or meta-analyses studies have been conducted on the seroprevalence of PPR in sheep and goats within South Asia, although similar studies have been reported in Ethiopia [9] and other parts of the world [5].

## 2 Materials and Methods

### 2.1 Guidelines and Protocol

Our research question, framed using the PECO framework (population, exposure, control, and outcome), focused on small ruminants (sheep and goats) as the population, the presence of PPRV in South Asia as the exposure, and the seroprevalence of pathogens as the outcome [24]. The study adhered to the Preferred Reporting Items for Systematic Reviews and Meta-Analyses (PRISMA) guidelines for conducting a systematic review and meta-analysis [25].

### 2.2 Systematic Literature Search Strategy

A comprehensive literature review was performed for this systematic review and meta-analysis to identify peer-reviewed studies reporting data on the seroprevalence of PPR in small ruminants (sheep and goats) from South Asian countries. Four authors (RC, MIH, MAR, and PB) rigorously investigated PubMed, Google Scholar, OpenAlex, and Science Direct for relevant peer-reviewed articles reporting primary research conducted from January 1, 2000, to June 1, 2025. A Boolean search strategy was employed, utilizing terms associated with PPR, seroprevalence, sheep, goats, small ruminants, and South Asian countries. The database search was conducted with filters based on the inclusion and exclusion criteria to ensure relevance.

### 2.3 Inclusion and Exclusion Criteria

For this systematic review and meta-analysis, specific inclusion and exclusion criteria were established to ensure the selection of relevant high-quality studies on the seroprevalence of PPR in sheep and goats within South Asian countries. Studies were included if they met the following criteria: (i) were peer-reviewed and published in English; (ii) reported primary research conducted between January 1, 2000, and June 1, 2025; (iii) were retrieved from recognized databases such as PubMed, Google Scholar, OpenAlex, and ScienceDirect; and (iv) focused specifically on the seroprevalence of PPR. The studies were excluded if they (i) did not report seroprevalence data, (ii) failed to mention the year of sample collection, (iii) were conducted outside South Asian regions, or (iv) involved blood samples collected from host species other than sheep and goats.

### 2.4 Data Collection and Extraction

We independently and meticulously conducted each stage of the review process, including the screening of titles and abstracts, full-text evaluation, data extraction, and quality assessment. The screening process commenced with the review of titles and abstracts to identify potentially eligible studies, followed by detailed full-text assessments. Studies were included if they met all predefined inclusion criteria (Tables 1-3). The articles were excluded if any essential information was missing or if they failed to meet the eligibility requirements.

**Table 1.** Extracted data from sheep and goats (combined)

| **Study** | **Total** | **Events** | **Country** | **Year** | **Vaccination** |
| --- | --- | --- | --- | --- | --- |
| [26] | 200 | 52 | Bangladesh | 2003 | No |
| [27] | 1463 | 1096 | Pakistan | 2005 | No |
| [28] | 660 | 286 | Pakistan | 2006 | No |
| [29] | 2798 | 1273 | Pakistan | 2006 | No |
| [30] | 1353 | 337 | Pakistan | 2006 | No |
| [31] | 3560 | 1339 | India | 2007 | No |
| [18] | 1056 | 510 | Pakistan | 2007 | No |
| [32] | 4884 | 2140 | India | 2009 | No |
| [33] | 389 | 77 | India | 2010 | No |
| [34] | 623 | 344 | Pakistan | 2010 | No |
| [35] | 461 | 190 | India | 2011 | No |
| [36] | 800 | 420 | Pakistan | 2011 | Yes |
| [37] | 70 | 24 | Pakistan | 2011 | No |
| [38] | 7096 | 2500 | Pakistan | 2013 | No |
| [39] | 391 | 70 | India | 2014 | No |
| [40] | 50 | 22 | Bangladesh | 2016 | No |
| [41] | 506 | 245 | India | 2016 | No |
| [42] | 460 | 380 | Nepal | 2016 | No |
| [43] | 246 | 144 | Pakistan | 2016 | No |
| [44] | 2420 | 1074 | Bangladesh | 2017 | No |
| [45] | 124 | 100 | Bangladesh | 2017 | No |
| [46] | 750 | 468 | India | 2017 | No |
| [47] | 398 | 260 | India | 2017 | No |
| [48] | 1000 | 450 | Pakistan | 2017 | No |
| [49] | 392 | 5 | India | 2018 | No |
| [50] | 776 | 9 | India | 2018 | No |
| [51] | 1836 | 844 | India | 2018 | No |
| [52] | 6643 | 3917 | India | 2018 | Yes |
| [53] | 5843 | 2574 | India | 2018 | Yes |
| [54] | 4687 | 2717 | India | 2018 | Yes |
| [55] | 598 | 43 | India | 2018 | No |
| [56] | 221 | 64 | Nepal | 2019 | No |
| [57] | 209 | 106 | India | 2021 | No |
| [58] | 5700 | 1587 | Pakistan | 2021 | No |
| [59] | 652 | 339 | Pakistan | 2021 | No |
| [60] | 80 | 26 | Bangladesh | 2022 | No |
| [61] | 1102 | 809 | India | 2023 | Yes |

**Table 2.**
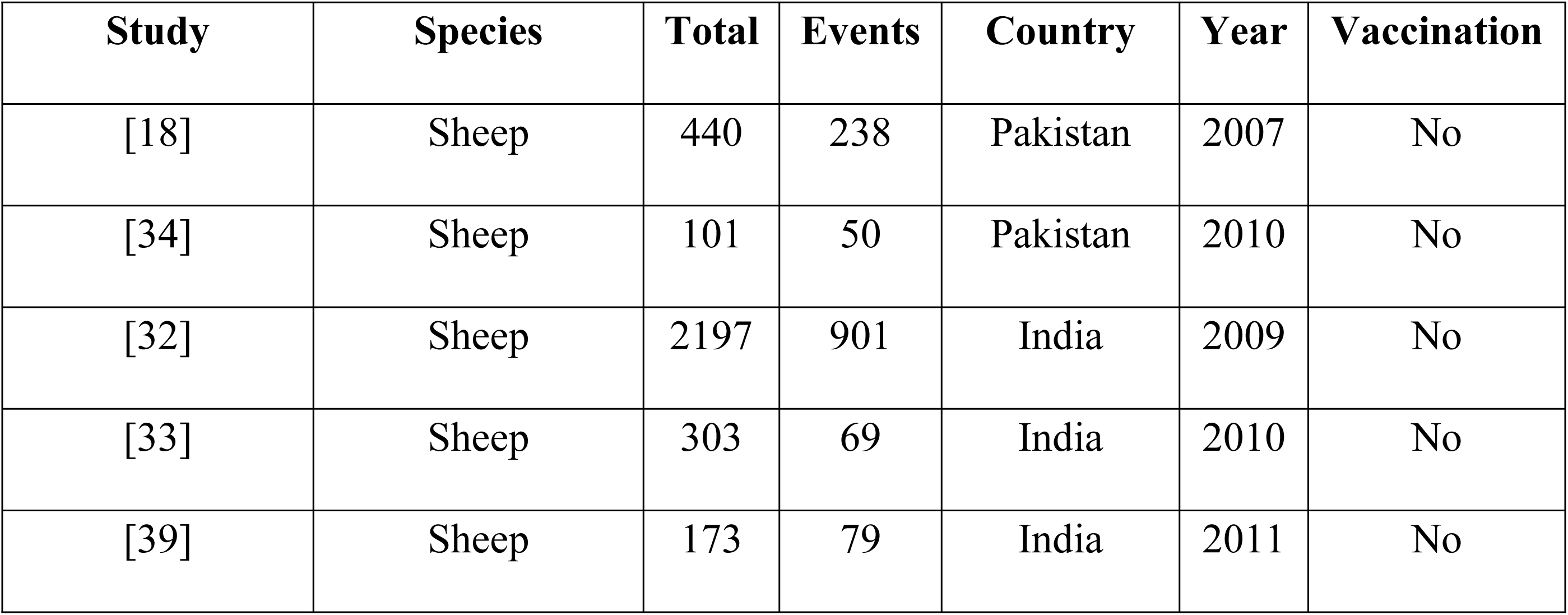
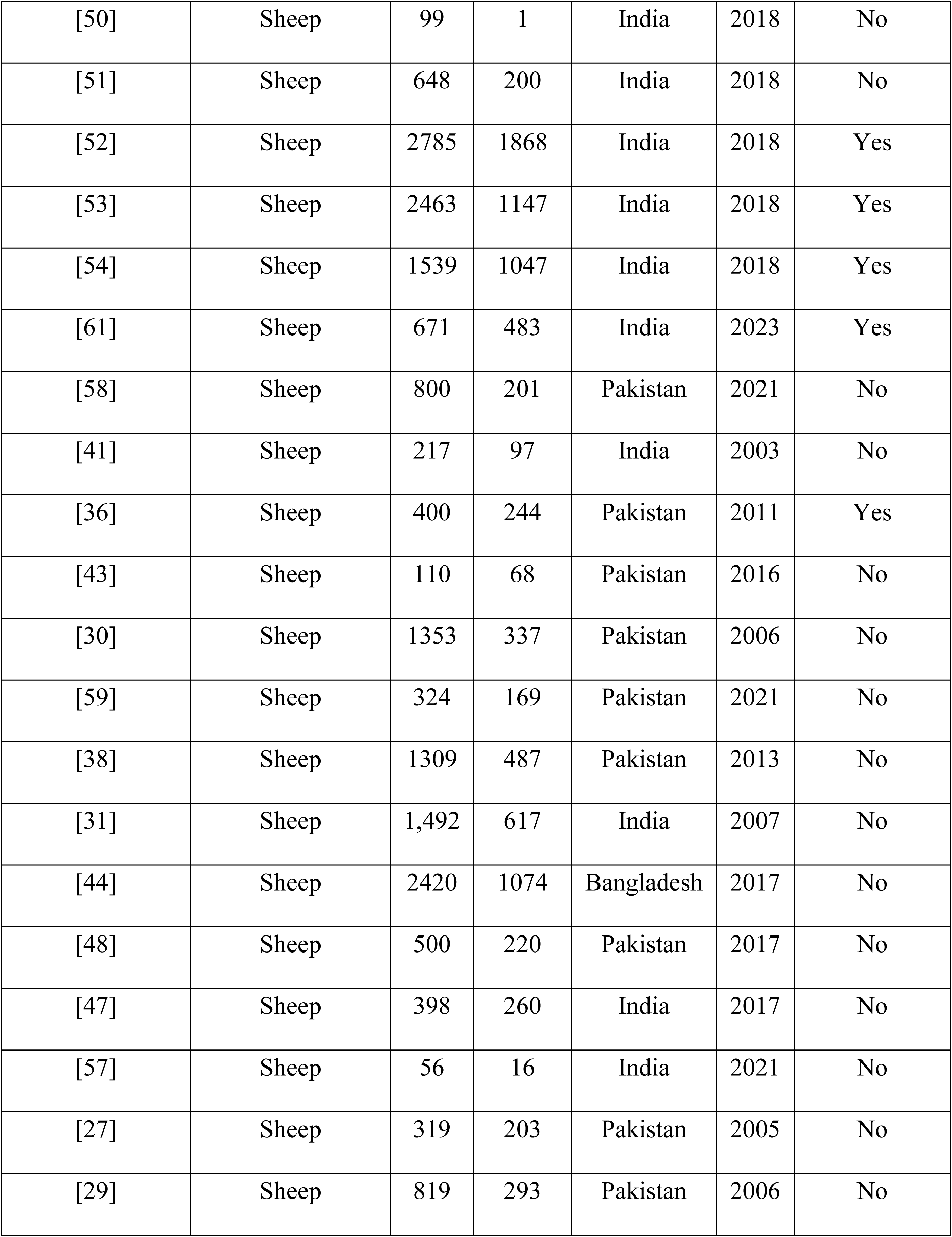
Extraction of data only from sheep.

| Study | Species | Total | Events | Country | Year | Vaccination |
| --- | --- | --- | --- | --- | --- | --- |
| [18] | Sheep | 440 | 238 | Pakistan | 2007 | No |
| [34] | Sheep | 101 | 50 | Pakistan | 2010 | No |
| [32] | Sheep | 2197 | 901 | India | 2009 | No |
| [33] | Sheep | 303 | 69 | India | 2010 | No |
| [39] | Sheep | 173 | 79 | India | 2011 | No |
| [50] | Sheep | 99 | 1 | India | 2018 | No |
| [51] | Sheep | 648 | 200 | India | 2018 | No |
| [52] | Sheep | 2785 | 1868 | India | 2018 | Yes |
| [53] | Sheep | 2463 | 1147 | India | 2018 | Yes |
| [54] | Sheep | 1539 | 1047 | India | 2018 | Yes |
| [61] | Sheep | 671 | 483 | India | 2023 | Yes |
| [58] | Sheep | 800 | 201 | Pakistan | 2021 | No |
| [41] | Sheep | 217 | 97 | India | 2003 | No |
| [36] | Sheep | 400 | 244 | Pakistan | 2011 | Yes |
| [43] | Sheep | 110 | 68 | Pakistan | 2016 | No |
| [30] | Sheep | 1353 | 337 | Pakistan | 2006 | No |
| [59] | Sheep | 324 | 169 | Pakistan | 2021 | No |
| [38] | Sheep | 1309 | 487 | Pakistan | 2013 | No |
| [31] | Sheep | 1,492 | 617 | India | 2007 | No |
| [44] | Sheep | 2420 | 1074 | Bangladesh | 2017 | No |
| [48] | Sheep | 500 | 220 | Pakistan | 2017 | No |
| [47] | Sheep | 398 | 260 | India | 2017 | No |
| [57] | Sheep | 56 | 16 | India | 2021 | No |
| [27] | Sheep | 319 | 203 | Pakistan | 2005 | No |
| [29] | Sheep | 819 | 293 | Pakistan | 2006 | No |

**Table 3.** Extraction of data from only goats.

| Study | Species | Total | Events | Country | Year | Vaccination |
| --- | --- | --- | --- | --- | --- | --- |
| [18] | Goat | 616 | 272 | Pakistan | 2007 | No |
| [34] | Goat | 522 | 294 | Pakistan | 2010 | No |
| [42] | Goat | 460 | 380 | Nepal | 2016 | No |
| [40] | Goat | 50 | 22 | Bangladesh | 2016 | No |
| [32] | Goat | 2687 | 1239 | India | 2009 | No |
| [33] | Goat | 86 | 8 | India | 2010 | No |
| [39] | Goat | 288 | 111 | India | 2011 | No |
| [35] | Goat | 391 | 70 | India | 2014 | No |
| [49] | Goat | 392 | 5 | India | 2018 | No |
| [50] | Goat | 677 | 8 | India | 2018 | No |
| [51] | Goat | 1188 | 644 | India | 2018 | No |
| [52] | Goat | 3858 | 2049 | India | 2018 | Yes |
| [53] | Goat | 3380 | 1427 | India | 2018 | Yes |
| [54] | Goat | 3148 | 1670 | India | 2018 | Yes |
| [61] | Goat | 431 | 326 | India | 2023 | Yes |
| [58] | Goat | 4900 | 1386 | Pakistan | 2021 | No |
| [26] | Goat | 100 | 25 | Bangladesh | 2003 | No |
| [41] | Goat | 289 | 148 | India | 2016 | No |
| [36] | Goat | 400 | 176 | Pakistan | 2011 | No |
| [43] | Goat | 136 | 76 | Pakistan | 2016 | No |
| [55] | Goat | 598 | 43 | India | 2018 | No |
| [28] | Goat | 428 | 167 | Pakistan | 2006 | No |
| [59] | Goat | 328 | 170 | Pakistan | 2021 | No |
| [37] | Goat | 70 | 24 | Pakistan | 2011 | No |
| [38] | Goat | 5787 | 2013 | Pakistan | 2013 | No |
| [31] | Goat | 2,068 | 722 | India | 2007 | No |
| [48] | Goat | 500 | 230 | Pakistan | 2017 | No |
| [46] | Goat | 634 | 417 | India | 2017 | No |
| [56] | Goat | 221 | 64 | Nepal | 2019 | No |
| [60] | Goat | 80 | 26 | Bangladesh | 2022 | No |
| [57] | Goat | 153 | 90 | India | 2021 | No |
| [45] | Goat | 124 | 100 | Bangladesh | 2017 | No |
| [27] | Goat | 1144 | 893 | Pakistan | 2005 | No |
| [29] | Goat | 1979 | 980 | Pakistan | 2006 | No |

### 2.5 Bias Assessment and Quality Assessment

In this systematic review, the methodological quality and risk of bias of the included studies on the seroprevalence of PPR were assessed using the Joanna Briggs Institute (JBI) critical appraisal tool. These tools are specifically designed for different study types and focus on evaluating internal validity by identifying potential sources of bias related to study design, conduct and analysis [62]. The assessment covered eight key domains: appropriateness of the research design, clarity in describing the study setting, adequacy of population demographics, quality and representativeness of the sample data, procedures for sample transfer and storage, diagnostic methods used (e.g., ELISA or other serological assays), clarity in reporting seropositive results, and transparency in data sources. Each criterion was evaluated using a binary scoring system, where a score of ’1’ was assigned for a "yes" response (indicating compliance with the criterion), and ’0’ was assigned for a "no" response (indicating noncompliance). The total score for each study ranged from 0 to 8 (**S1 Table 1**). Based on these scores, the studies were categorized into three levels of bias risk: low risk of bias (total score >70%), moderate risk of bias (score between 50% and 69%), high risk of bias (score <50%). To ensure the robustness and reliability of the meta-analysis, sensitivity analyses were performed to assess the effect of excluding studies with moderate and high risk of bias on the overall pooled estimates. This structured and transparent approach to bias assessment contributes to the methodological rigor of the review by systematically identifying and mitigating the influence of potential biases in the included studies.

### 2.6 Statistical Analysis

The data that were retrieved and then entered into an Excel spreadsheet. As part of this comprehensive meta-analysis, the authors meticulously reviewed and assessed each article that met the inclusion criteria to ensure the integrity and reliability of the synthesized findings. By using a random effects model meta-analysis, the pooled seroprevalence of PPR at the 95% CI was determined [63,64]. Heterogeneity is classified as low, moderate, or high at I² values of 25%, 50%, and 75%, respectively, with 0% indicating its absence [65]. A random-effects meta-analysis was chosen for summary statistics because of the substantial heterogeneity across studies. A subgroup analysis was conducted by country and vaccination. Moreover, univariate meta-regression was conducted to explore potential sources of heterogeneity. The pooled seroprevalence of PPR is displayed on a map. Funnel plots and sensitivity analyses were employed for visual inspection of distribution, quality, and bias across articles. The significance threshold was set at p < 0.05. Meta-analysis of the data, including meta-regression, subgroup analysis, forest plots, funnel plots, and sensitivity analysis, was conducted via Stata version 18.0 (College Station, 163 TX, USA). R programming (version 4.5.0) was used to create the map using different packages such as “*ggplot”* and “*spdep*” [66,67]. This systematic review and meta-analysis were based on the PRISMA statement [68].

### 2.7 Ethics Statement

This study is a systematic review and meta-analysis and did not involve any direct experimentation on animals by the authors. Therefore, approval from an Institutional Animal Care and Use Committee (IACUC) or equivalent ethics board was not required. All data analyzed were obtained from previously published studies that stated appropriate ethical approval and animal care standards.

## 3. Results

### 3.1 Search Results and Eligible Studies

The PRISMA flow diagram illustrates the process of study selection for the systematic review and meta-analysis of the seroprevalence of PPR in sheep and goats across South Asia (Fig. 1). A total of 153 (n=153) records (reports may be better terms here) were identified through different database searches (PubMed, Google Scholar, OpenAlex, and ScienceDirect). After removing duplications (n = 47), 106 (n = 106) reports were screened based on their titles and abstracts. Among these, 19 (n=19) reports were excluded because they did not meet the eligibility criteria. The full texts of 53 (n=53) articles were assessed, whereas 34 (n=34) were excluded for reasons such as not reporting seroprevalence data, missing sampling years, and/or studies conducted outside South Asia. Ultimately, 37 (n=37) studies met the inclusion criteria and were included in the final analysis.

**Fig 1.**
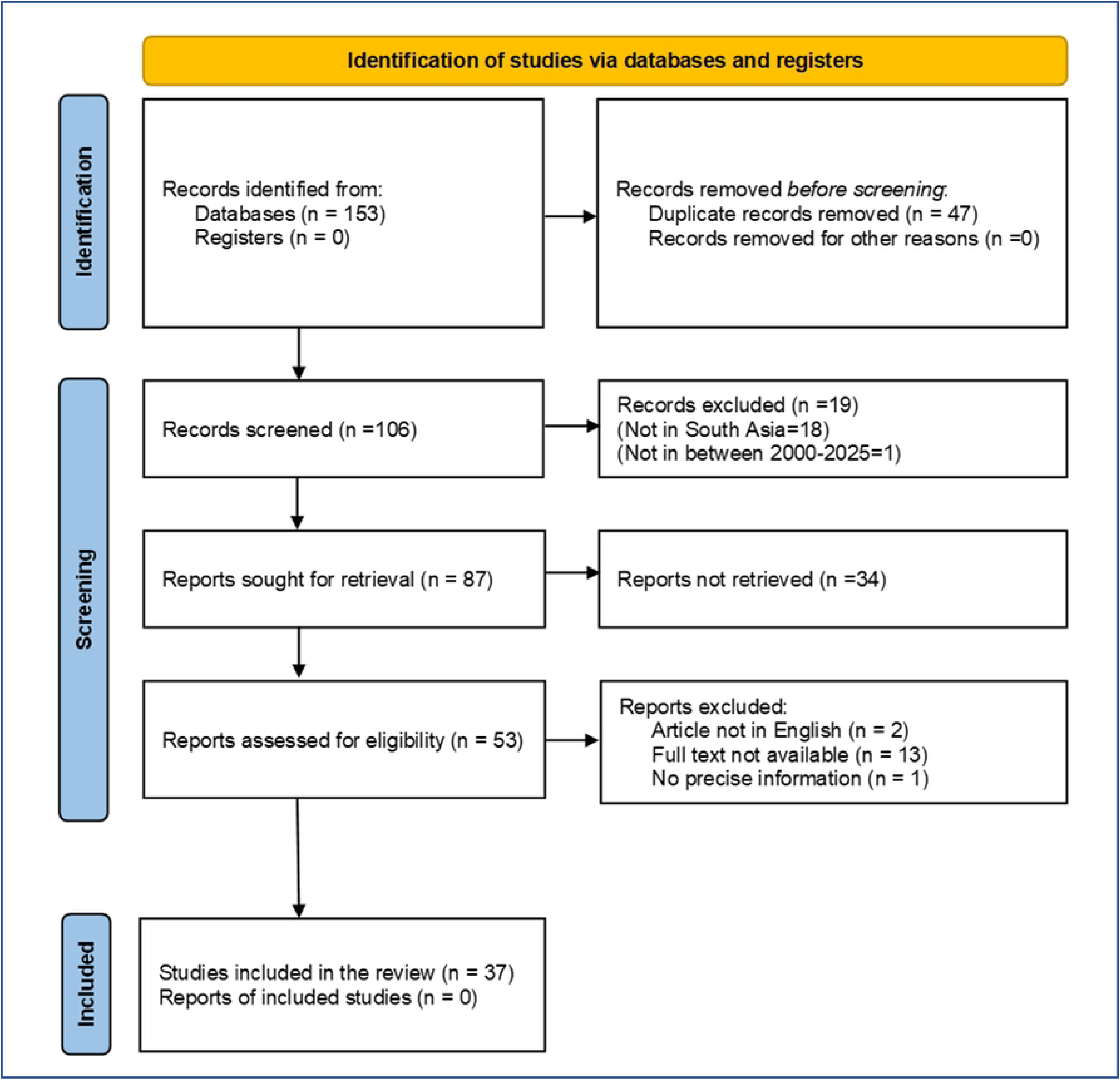
PRISMA flow diagram for the selection of studies.

### 3.2 Descriptive Characteristics of the Included Studies

The study included 37 (n=37) studies that met the eligible criteria for inclusion in this systematic review and meta-analysis from four (4) southern asian countries: Bangladesh (n=5) [26,40,44,45,60], India (n=17) [31–33,39,41,46,47,49–55,57,61,69], Pakistan (n=13) [18,27–30,34,36–38,43,48,58,59], and Nepal (n=2) [56,70] (Fig. 2).

**Fig 2.**
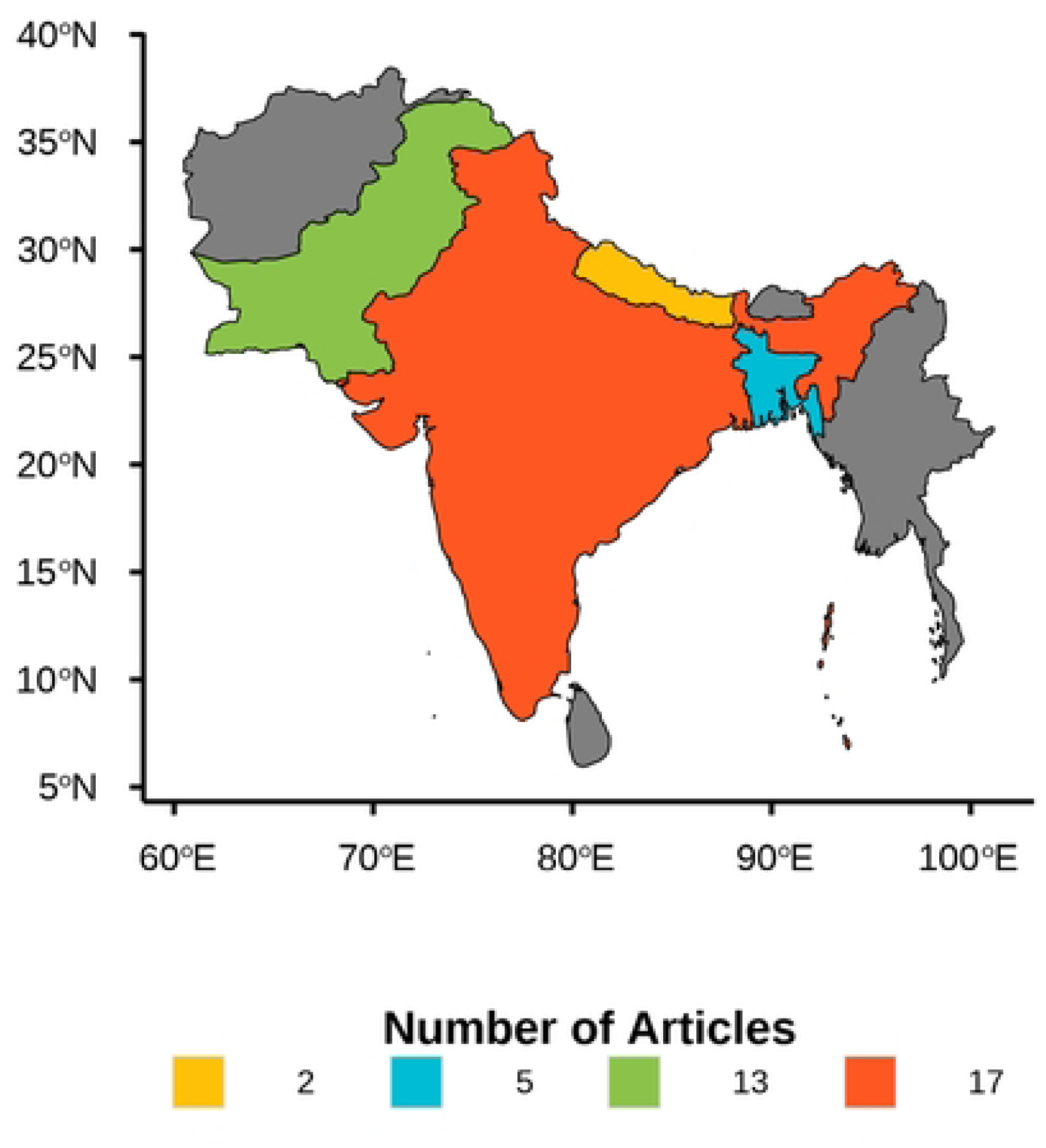
A map illustrating the geographical distribution of the selected studies for this meta-analysis (n = 37) was generated via the ggplot2 package in R software (v 4.5.0) [67].

### 3.3 Pooled seroprevalence of PPR

The forest plot illustrates the pooled prevalence of PPR in South Asia, as shown in Figs. 3-5. Even though there are some differences across the different studies, the heterogeneity is significant; hence, the random effects model was selected. The random effects meta-analysis showed that the overall pooled prevalence of PPR in sheep and goats was 42.4% (95% CI: 35.0–49.9%) with considerable heterogeneity (I^2^ = 99.7%, τ2 = 0.22, p<0.001) (Fig. 3). Moreover, the overall pooled prevalence of PPR in goats was 41.7% (95% CI: 33.7–50.1%) with heterogeneity (I^2^ = 99.6%, τ^2^ = 0.24, p<0.001) (Fig. 4). Furthermore, the estimated prevalence of PPR in sheep was 44.7% (95% CI: 37.0–52.0%) with heterogeneity (I^2^ = 99.2%, τ^2^ = 0.15, p<0.001) (Fig. 5). A geographical map was created to illustrate the prevalence of PPR in South Asian countries in this study (Fig. 6).

**Fig 3.**
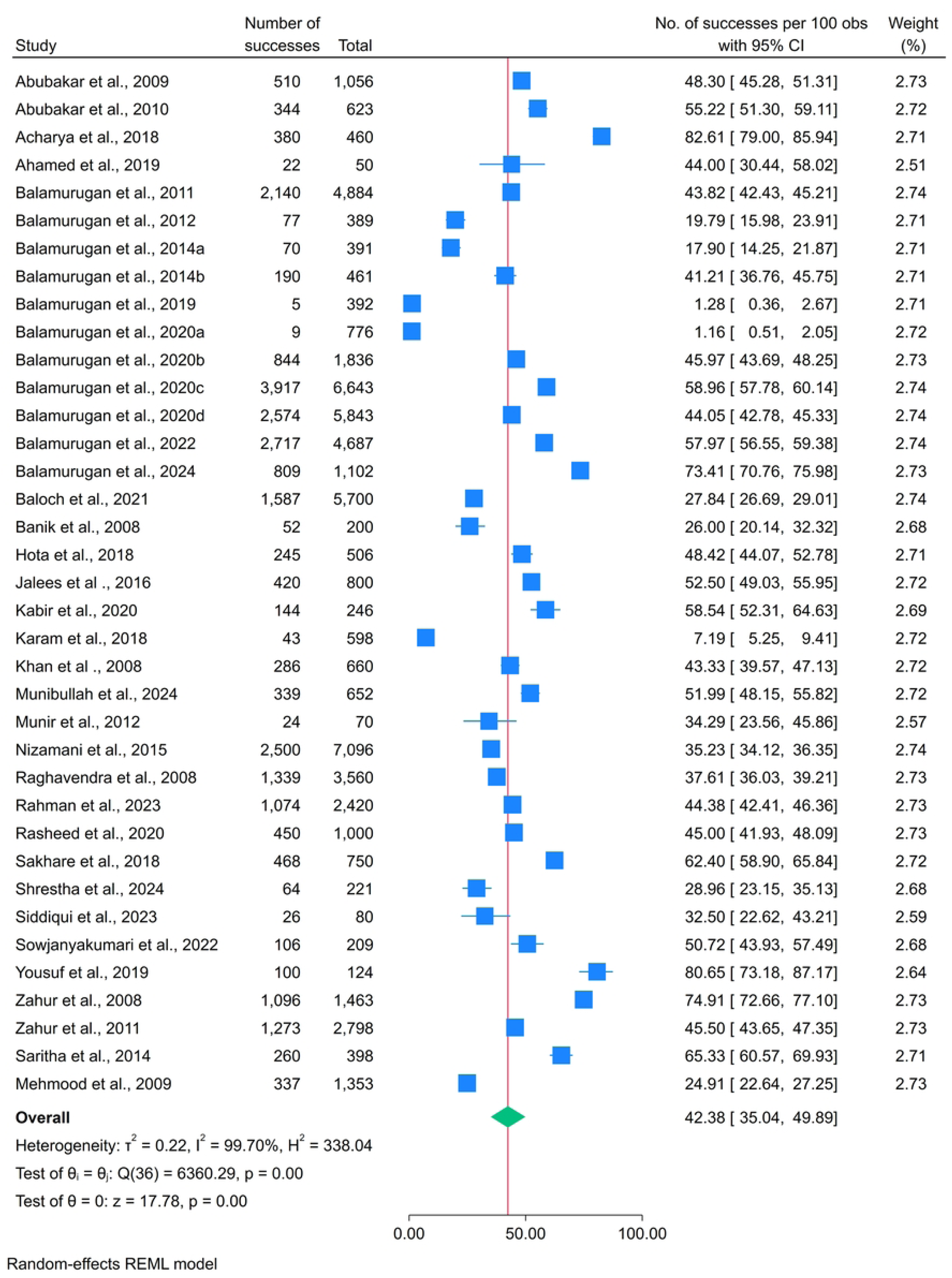
Forest plots for random-effects meta-analysis of PPR in sheep and goats.

**Fig 4.**
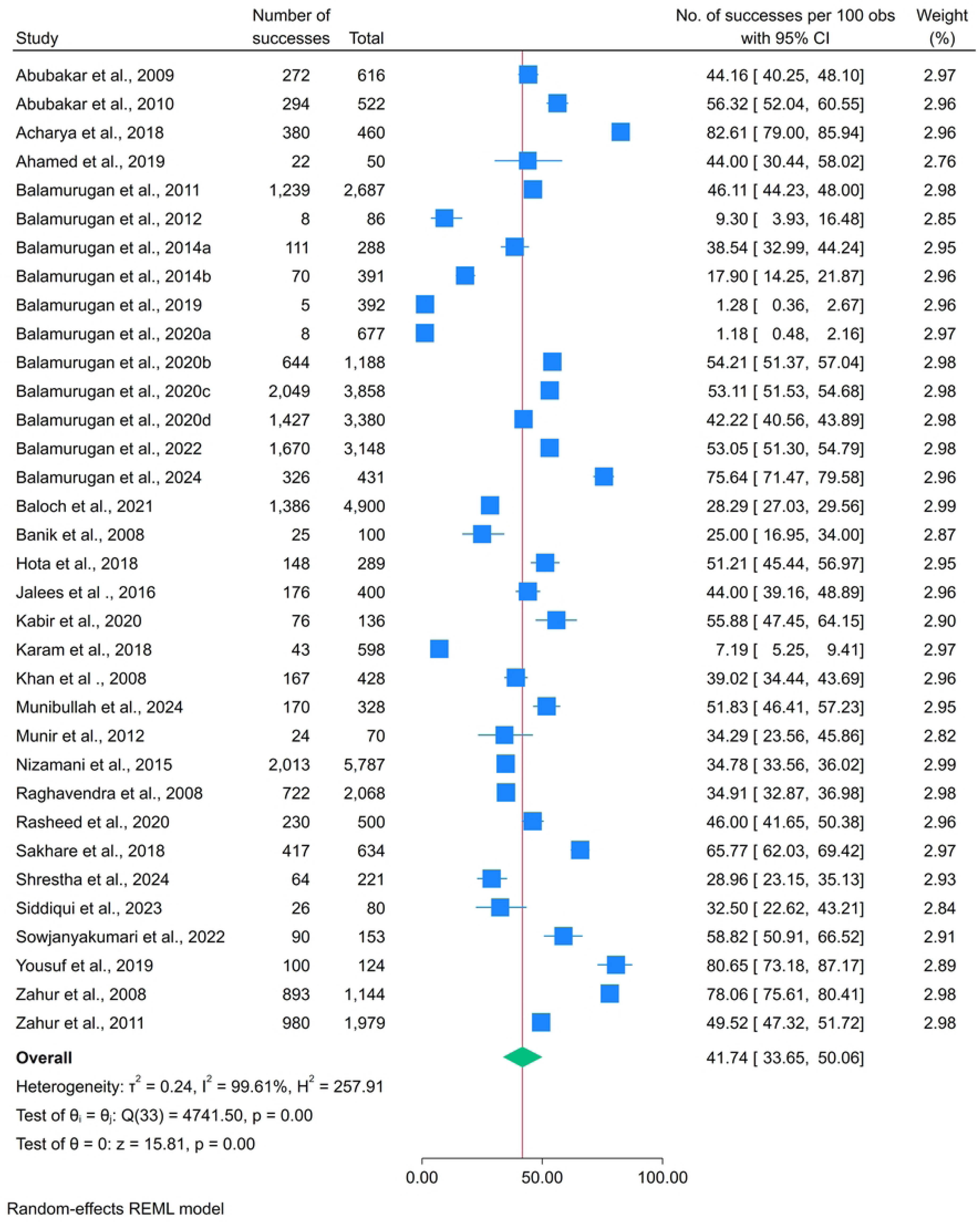
Forest plots for random-effects meta-analysis of PPR in goats.

**Fig 5.**
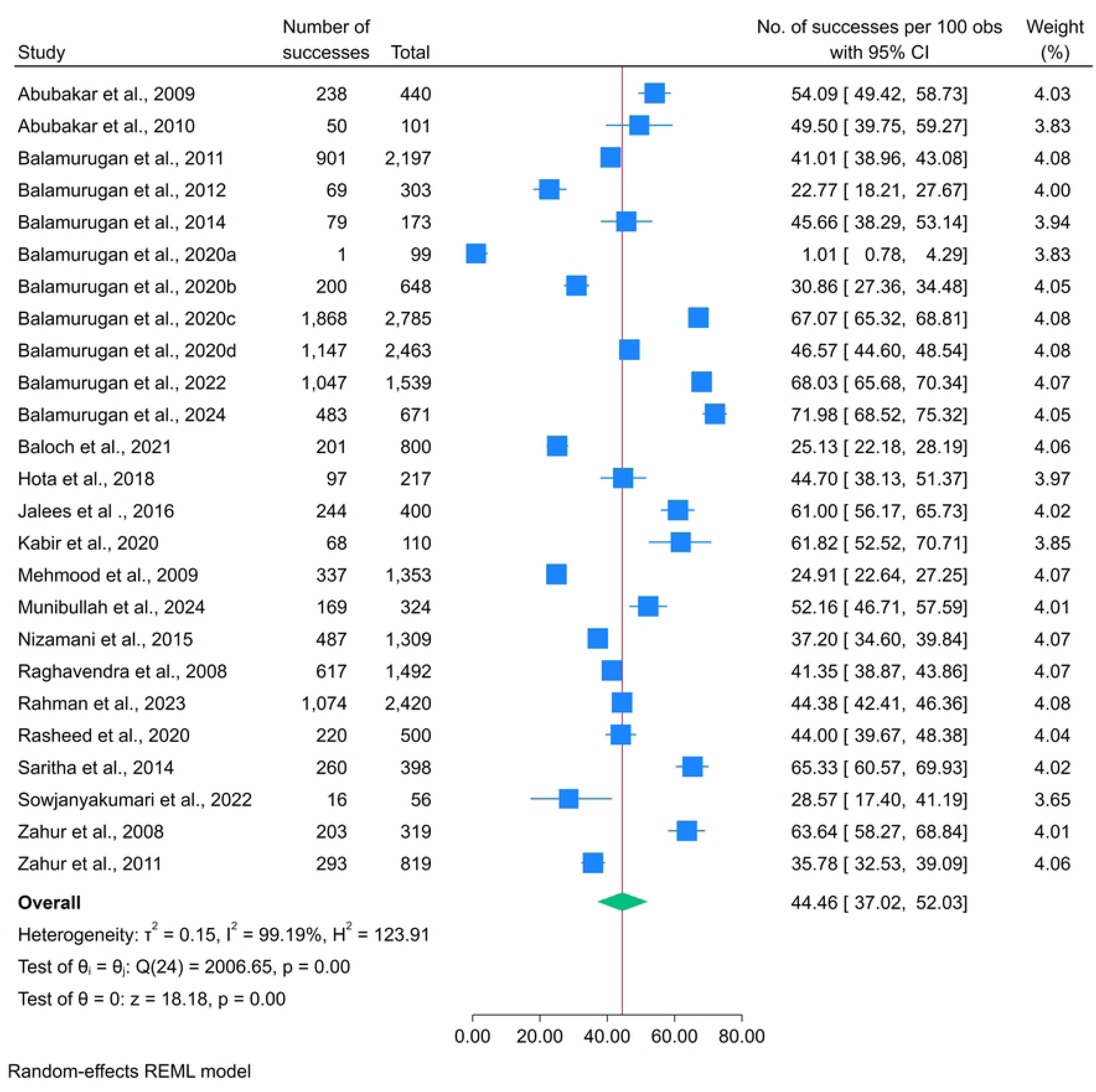
Forest plots for random-effects meta-analysis of PPR in sheep.

**Fig 6.**
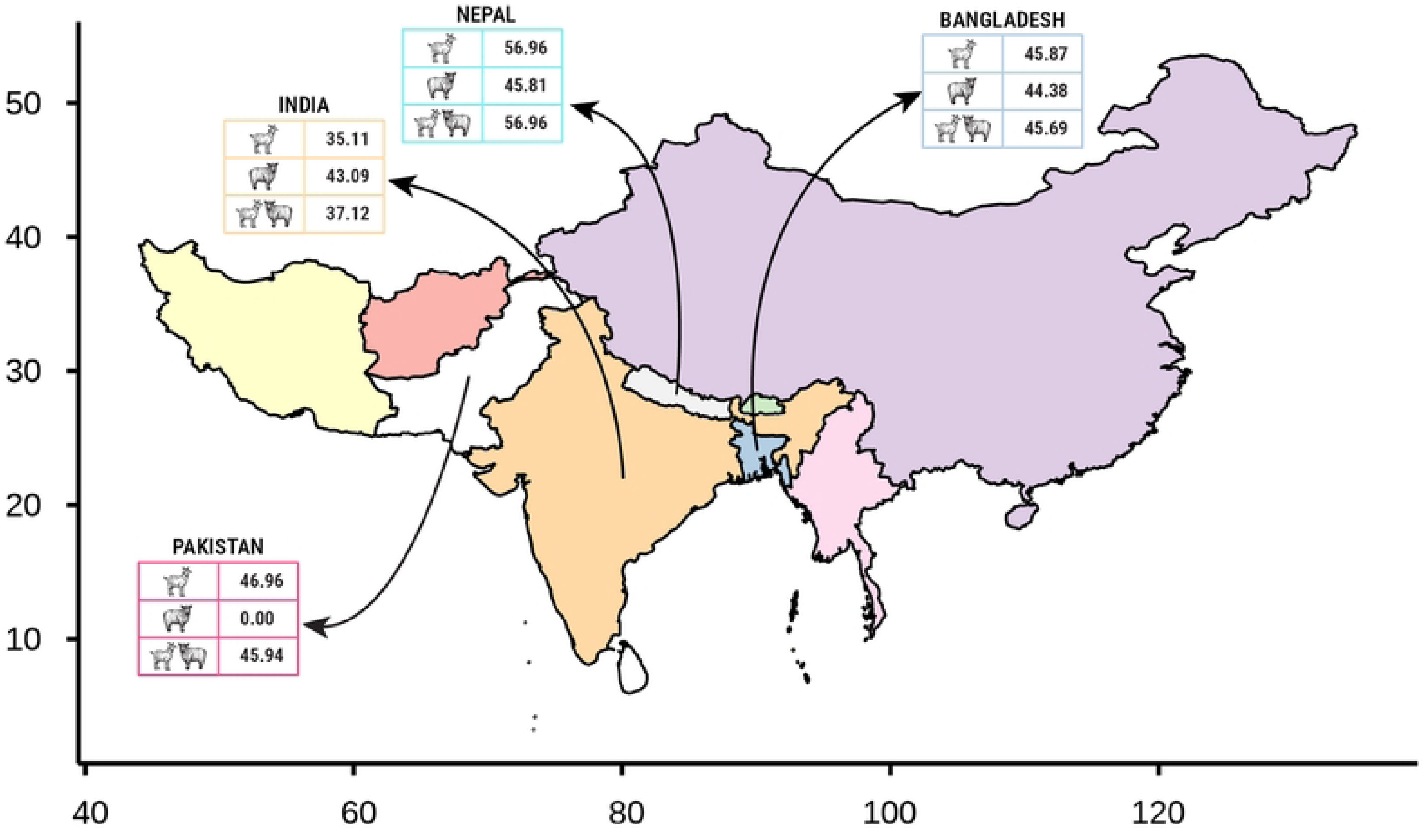
The prevalence of Peste des Petits Ruminants (PPR) among sheep, goats, and total sheep and goats in South Asia is illustrated in Figures 3–5. These maps were generated using R programming to spatially represent the distribution patterns.

### 3.4 Subgroup pooled prevalence of PPR

A subgroup meta-analysis was performed for the country and vaccination status (Figs. 7-12). For both sheep and goats, the pooled prevalence of PPR of the country was 45.7% (Bangladesh), 37.1% (India), 57.0% (Nepal), and 45.9% (Pakistan). The highest prevalence rate (57.0%) was reported in Nepal (95% CI: 8.4-97.7), and the lowest prevalence rate (37.1%) was reported in India (95% CI: 24.9-50.3) (Fig. 7). In terms of vaccination status, the pooled prevalence was greater for vaccinated sheep and goats, at 57.5% (95% CI: 47.9-66.9) (Fig. 8). For goats, the estimated pooled prevalence of PPR by country was 45.9%, 35.1%, 57.0%, and 47.0% in Bangladesh, India, Nepal, and Pakistan, respectively (Fig. 9). Vaccination status significantly influenced the prevalence, as the pooled prevalence was higher (56.2%) in vaccinated goat (95% CI: 42.0-69.8) than in unvaccinated goats 39.8% (Fig. 10). Moreover, the pooled prevalence of PPR in sheep among the countries included in this study was 44.4%, 43.1%, and 45.8% in Bangladesh, India, and Pakistan, respectively, but no study was conducted in Nepal (Fig. 11). In terms of vaccination status, vaccinated sheep were found to have a higher pooled prevalence of 63.1% (95% CI: 54.2-71.5) (Fig. 12).

**Fig 7.**
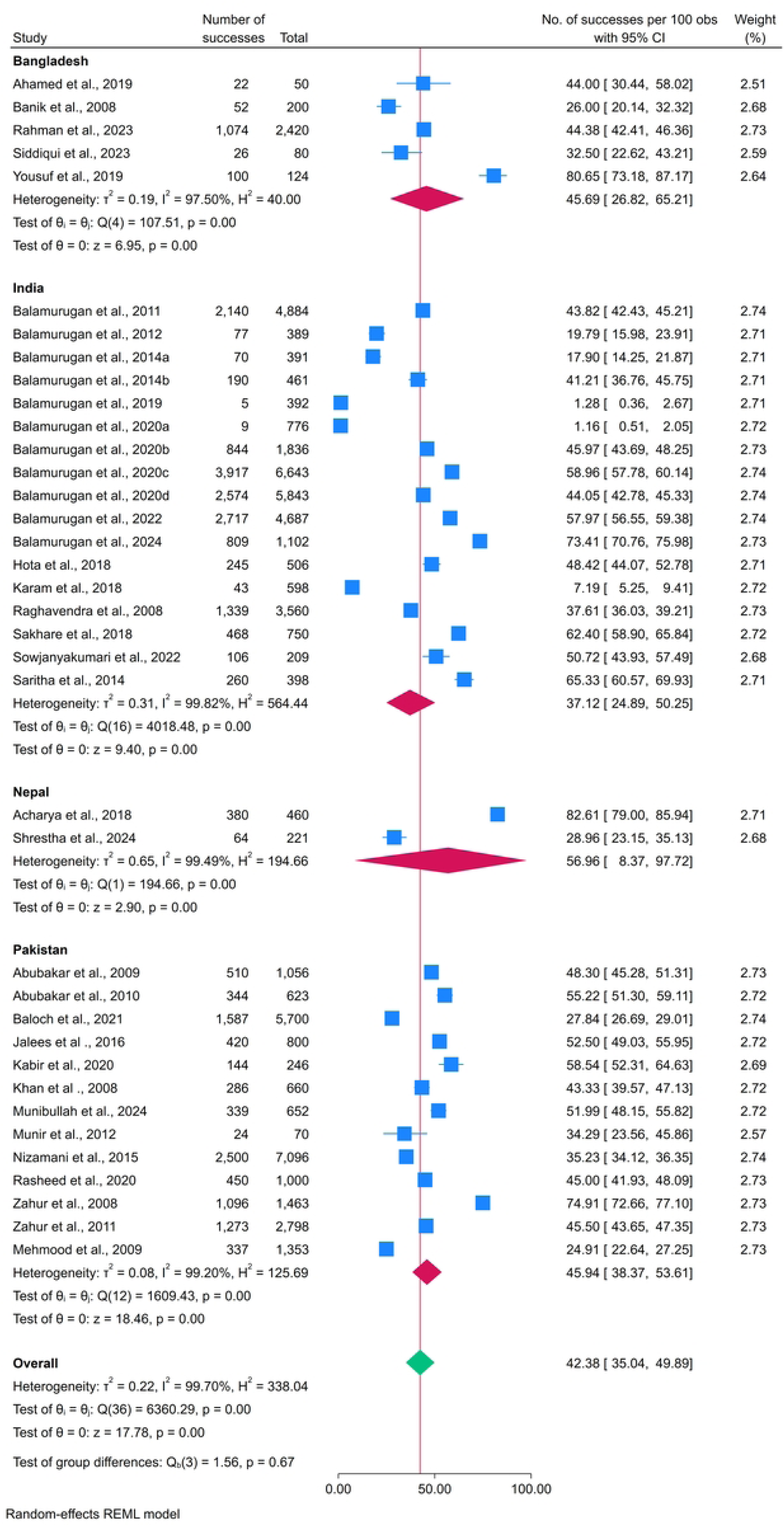
Subgroup Analysis by Country for PPR Prevalence in Sheep and Goats.

**Fig 8.**
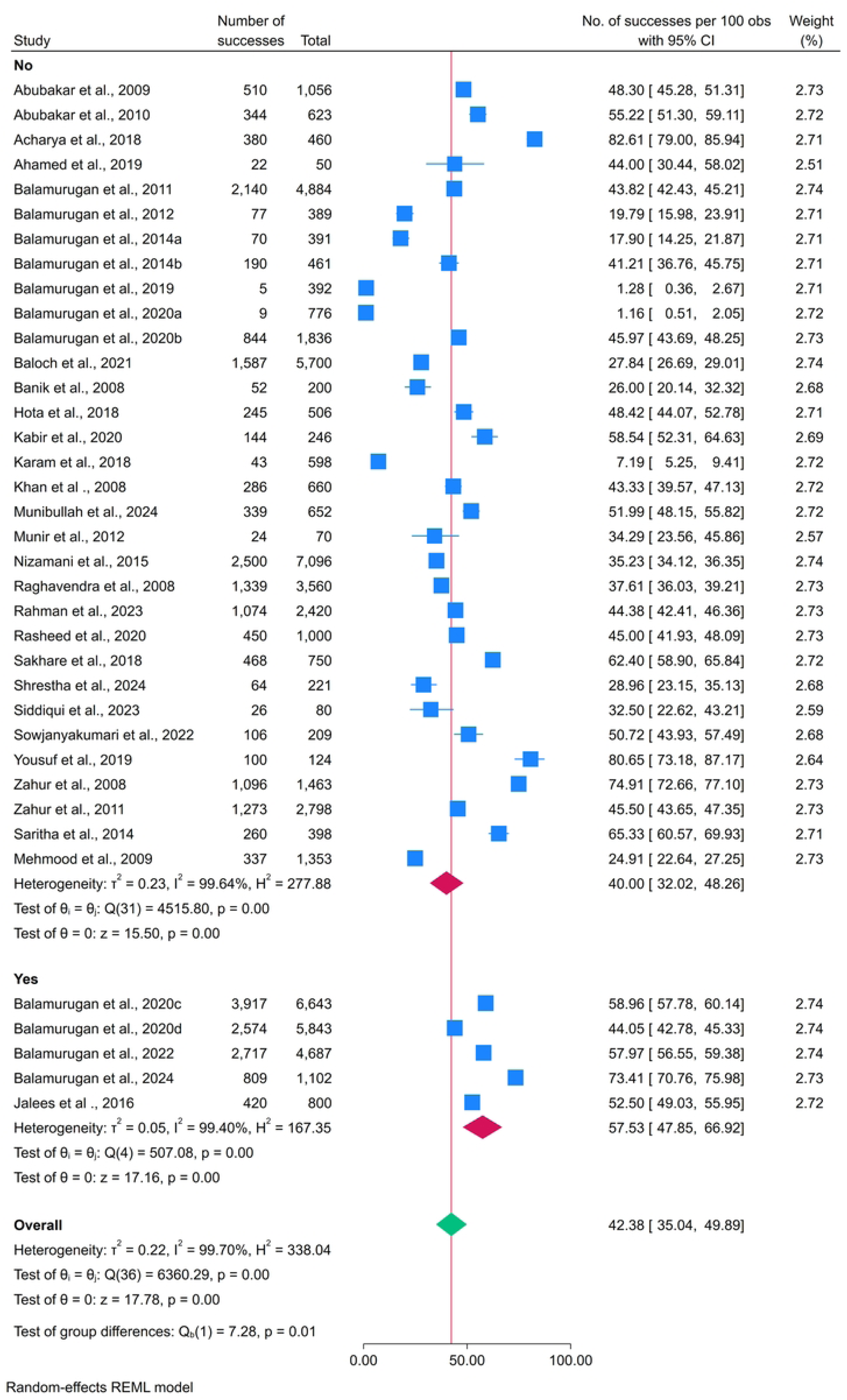
Subgroup Analysis by Vaccination for PPR Prevalence in Sheep and Goats.

**Fig 9.**
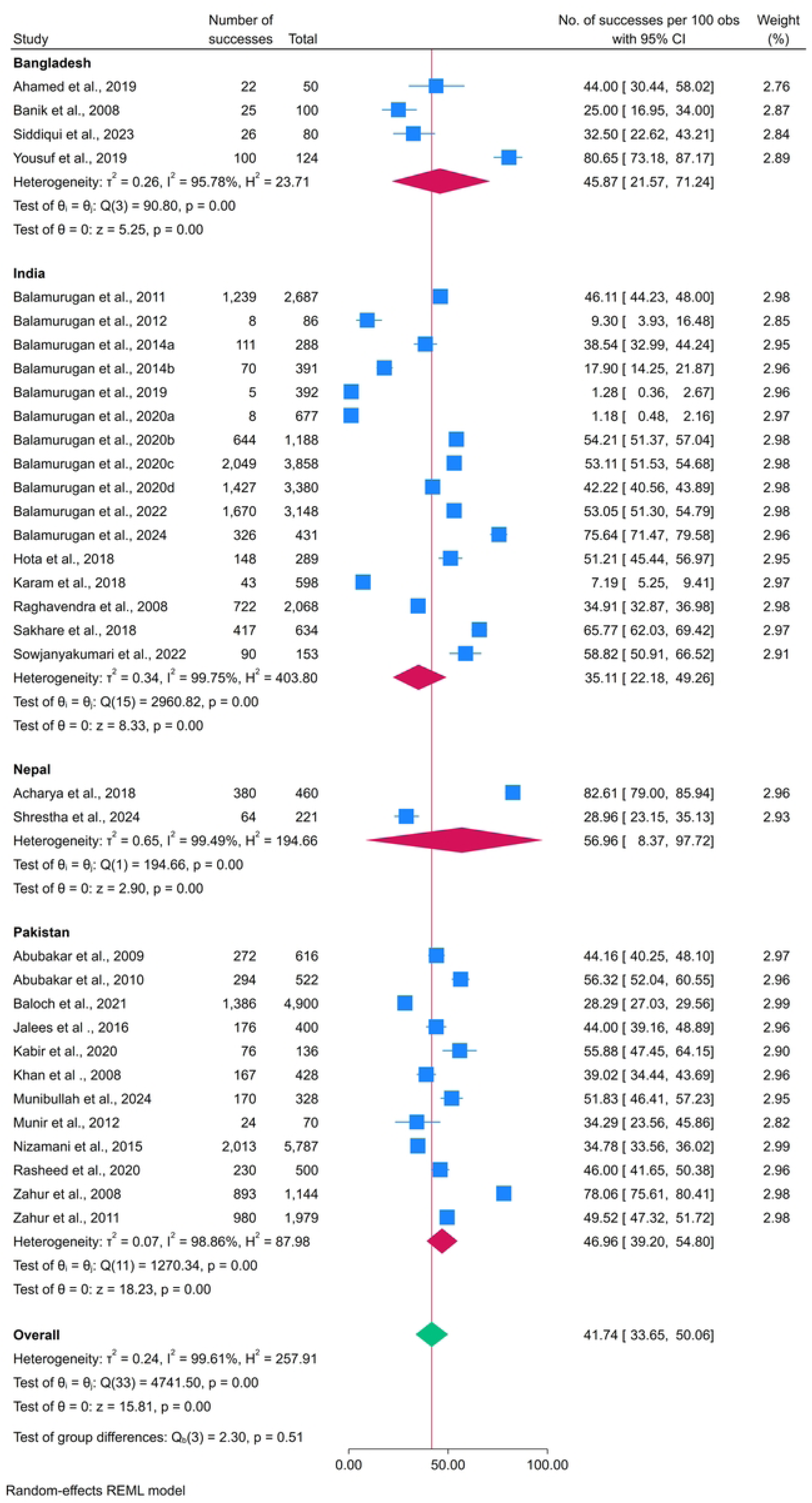
Subgroup Analysis by Country for PPR Prevalence in Goats.

**Fig 10.**
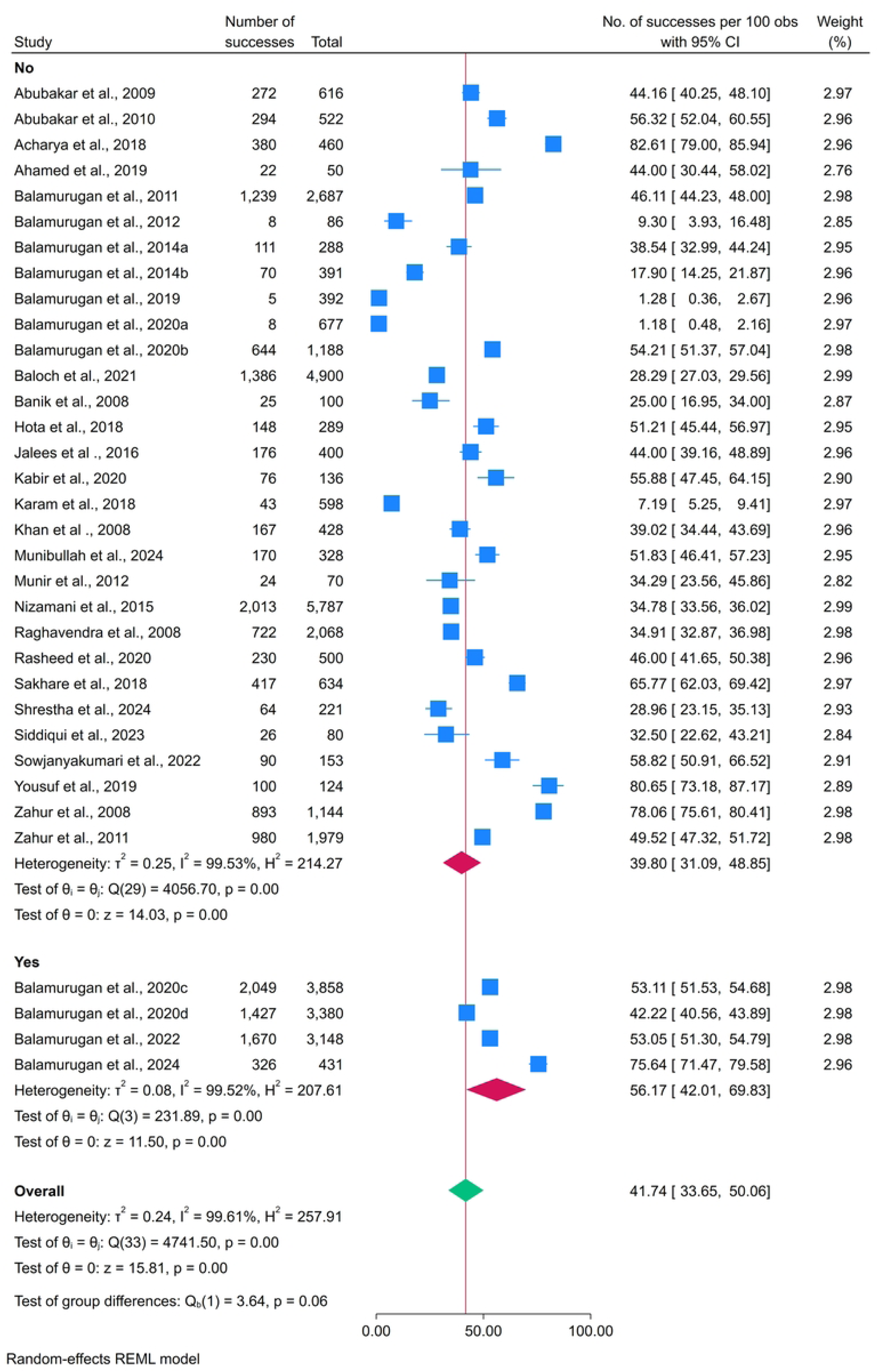
Subgroup Analysis by Vaccination for PPR Prevalence in Goats.

**Fig 11.**
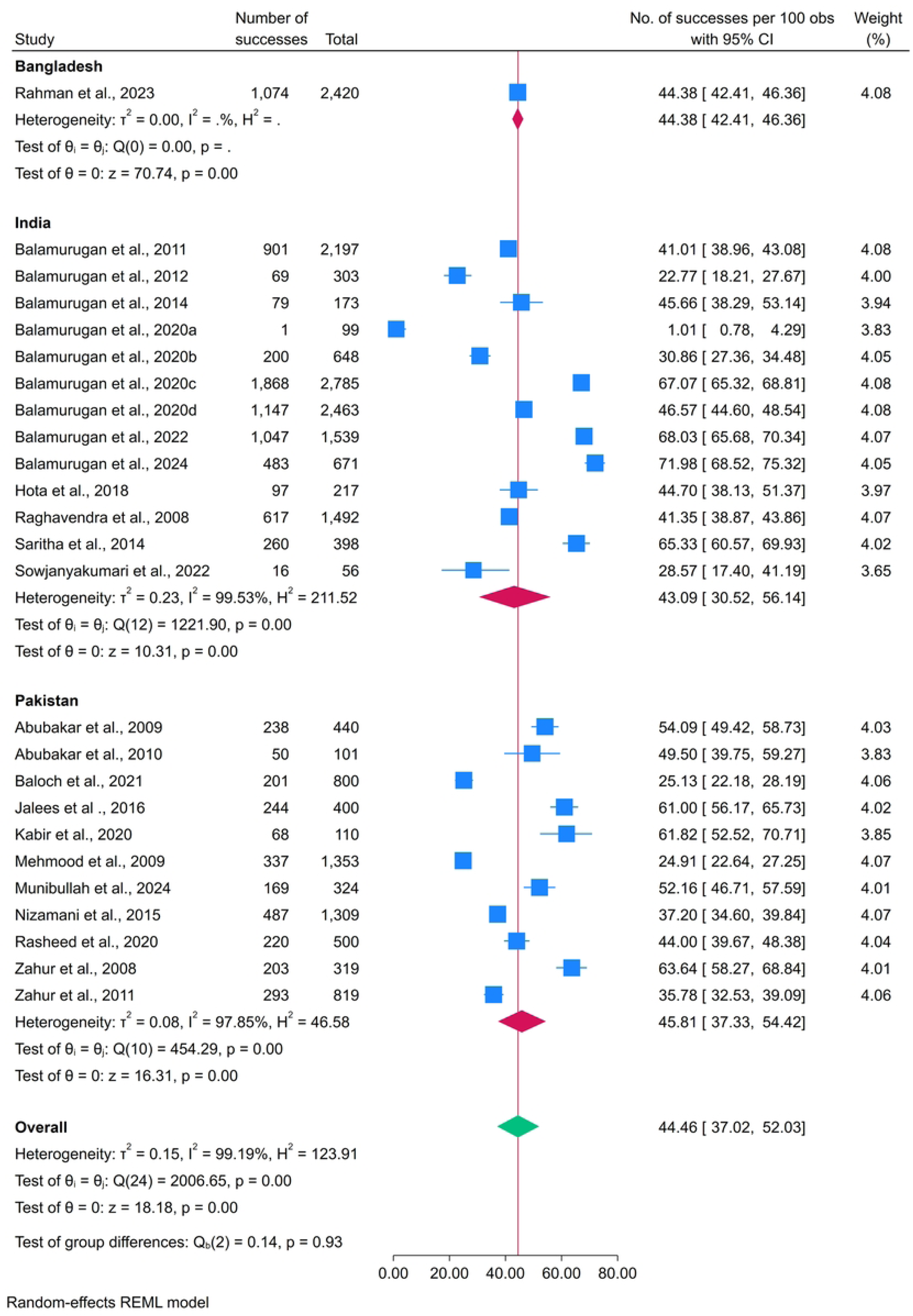
Subgroup Analysis by Country for PPR Prevalence in Sheep.

**Figure 12.**
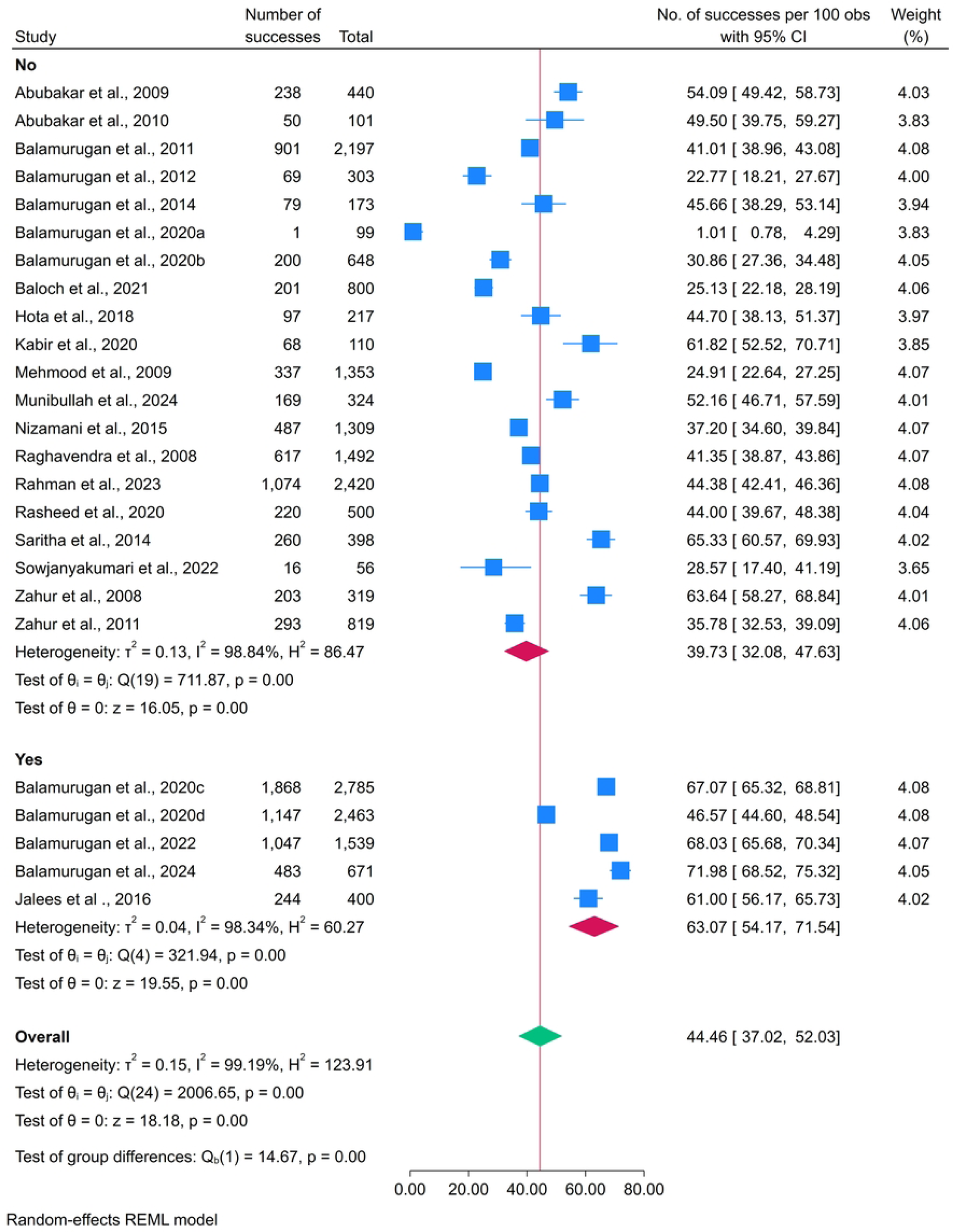
Subgroup Analysis by Vaccination for PPR Prevalence in Sheep.

### 3.5 Sensitivity analysis and meta-regression of covariants

A leave-one-out sensitivity analysis was performed to evaluate the influence of individual studies on the pooled prevalence estimation of PPR among sheep and goats, and only among sheep and goats. The analysis encompassed each study sequentially, excluding it to assess its impact on the overall estimation. The pooled prevalence remained stable, ranging from 0.41 (95% CI: 0.34–0.49) to 0.44 (95% CI: 0.37–0.51) in sheep and goats, 0.43 (95% CI: 0.36—0.51) to 0.47 (95% CI: 0.41—0.53) in sheep (only) and 0.40 (95% CI: 0.33—0.49) to 0.43 (95% CI: 0.35—0.51) in goats (only), demonstrating that no single study exerted a disproportionate effect on the summary estimation (Figs. 16-18). The funnel plots demonstrated a relatively symmetrical distribution of the studies around the pooled prevalence estimation, suggesting minimal evidence of publication bias (Figs. 13-15). Univariate meta-regression showed that vaccination status was significantly associated with PPR prevalence in sheep (β = 0.4704, p =0.005, R² = 23.39%), whereas other covariates, including study year, country, sample size, and positive cases, had no significant effect. No meaningful associations were observed for the goats or sheep, or combined sheep and goats data (Table 4).

**Fig 13.**
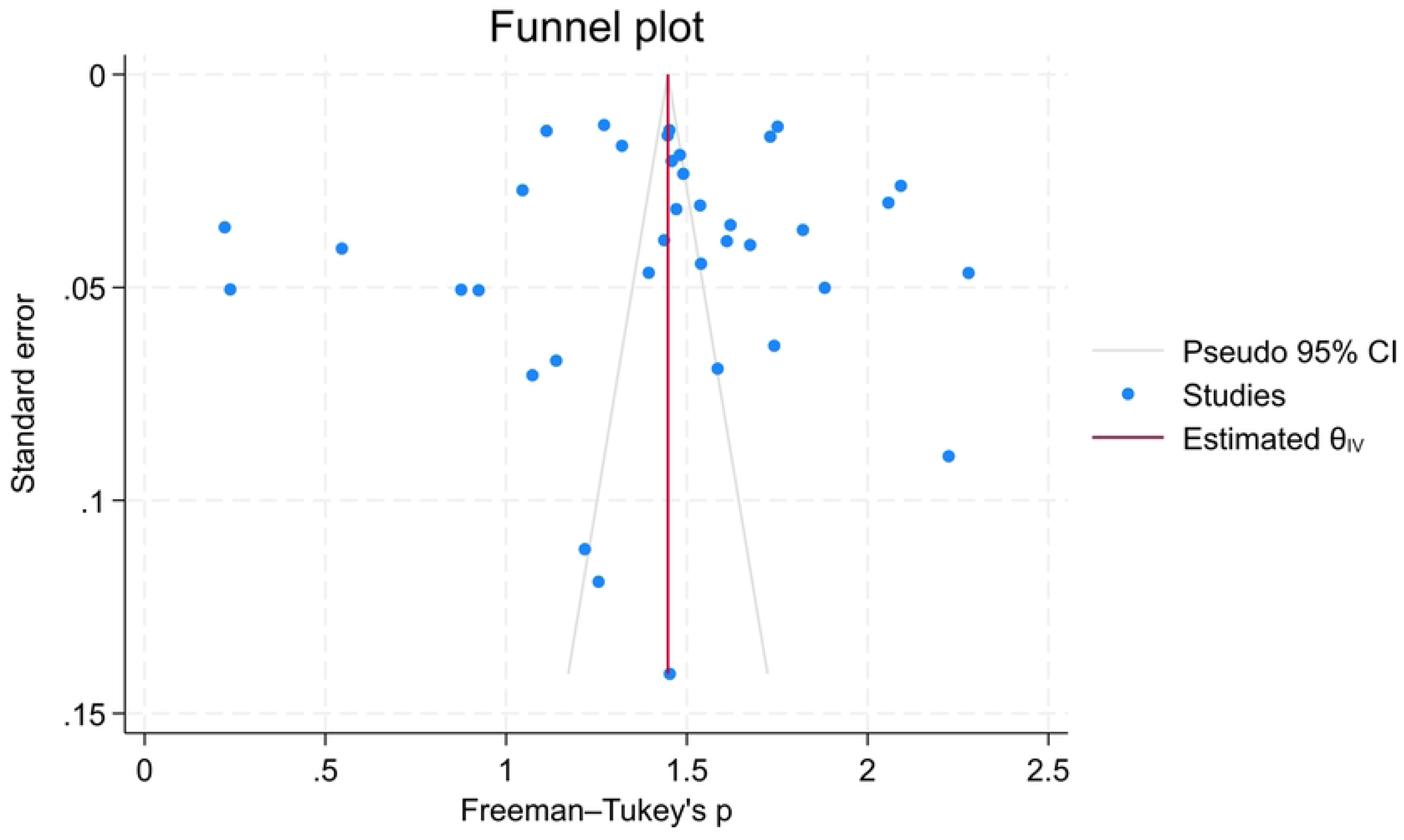
Funnel plot to assess publication bias in studies evaluating *PPR* in sheep and goats.

**Fig 14.**
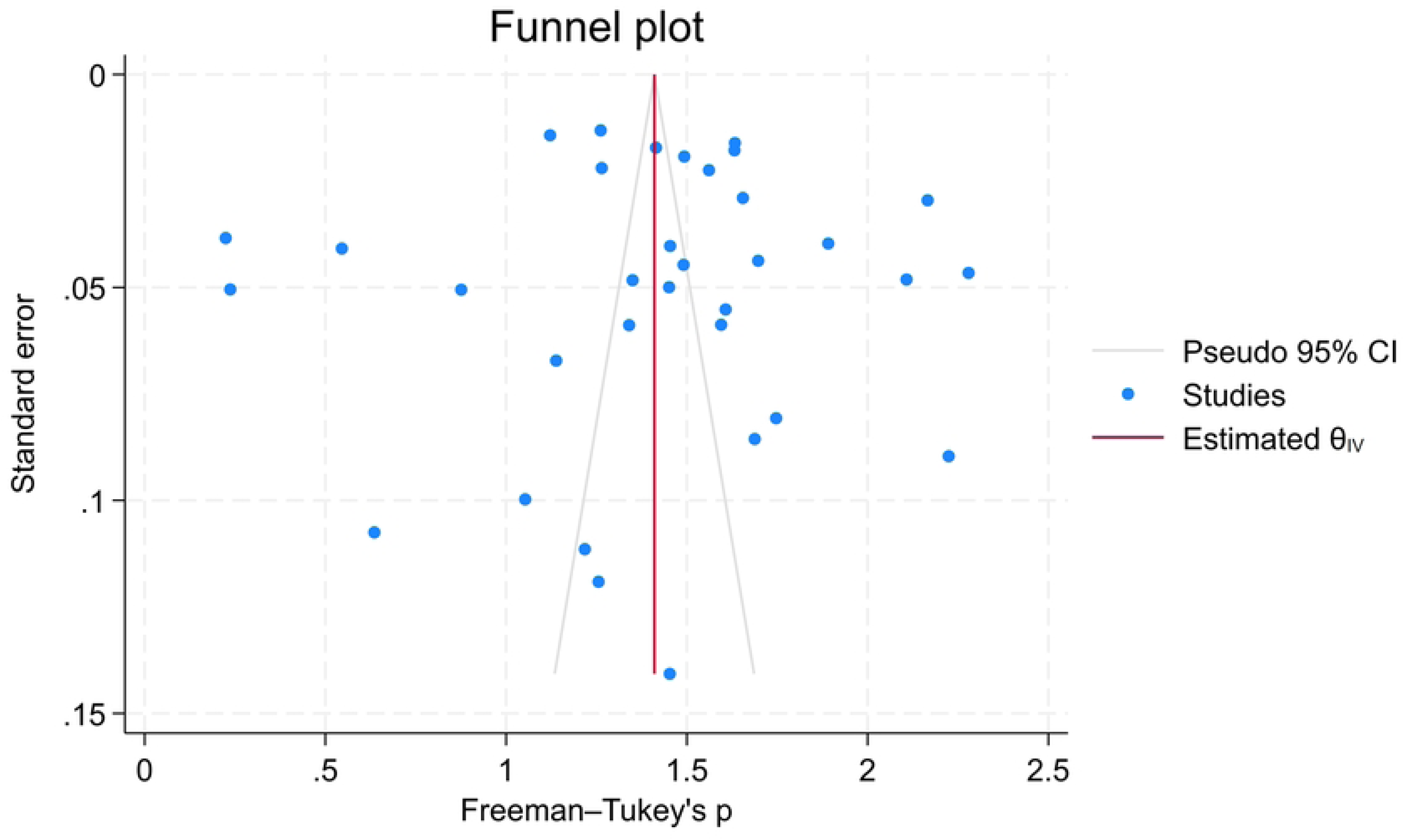
Funnel plot to assess publication bias in studies evaluating PPR in goats.

**Fig 15.**
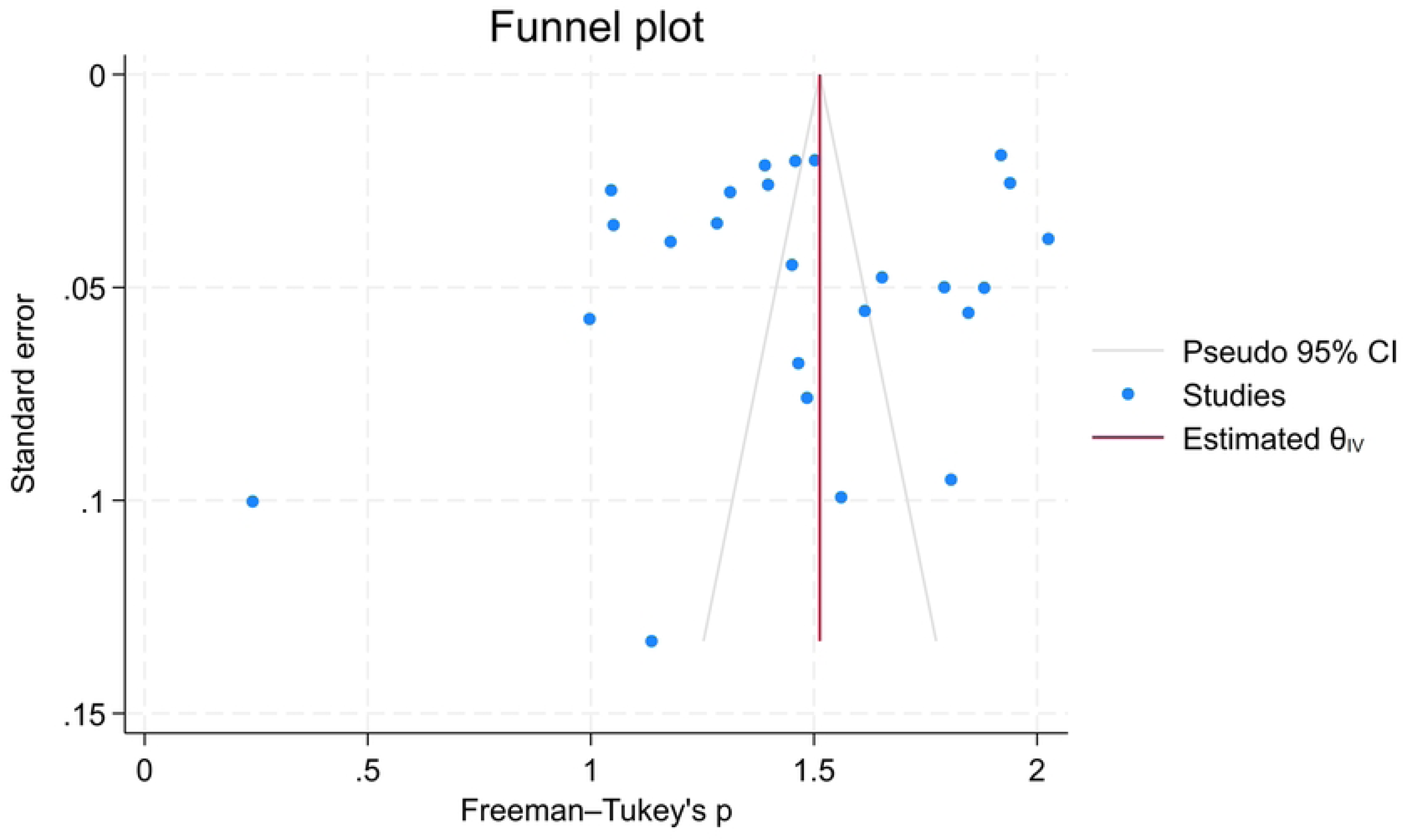
Funnel plot to assess publication bias in studies evaluating *PPR* in sheep.

**Fig 16:**
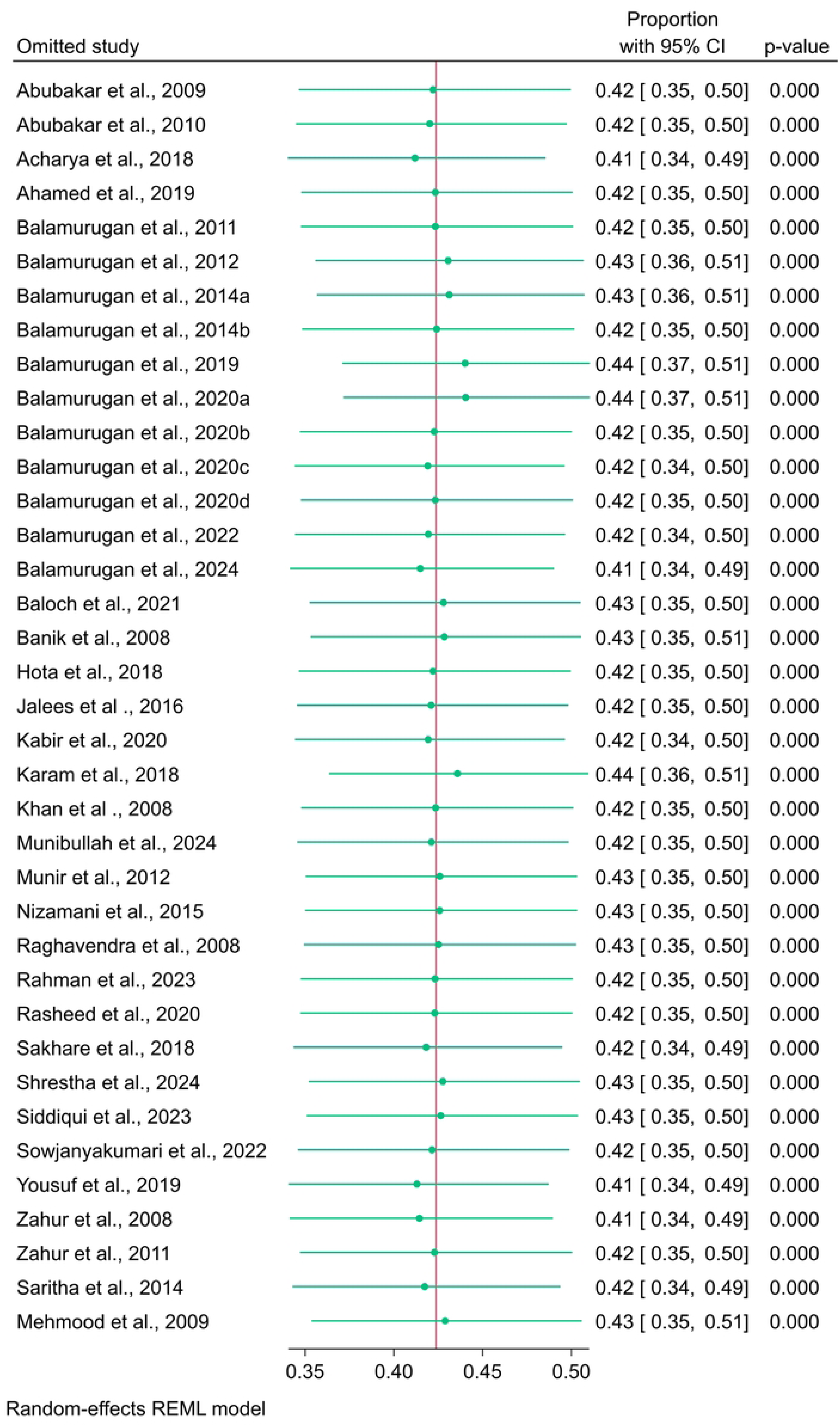
Leave-One-Out Sensitivity Analysis Plot for the PPR in Sheep and Goats. Each point represents the pooled prevalence estimate when the corresponding study is omitted.

**Fig 17:**
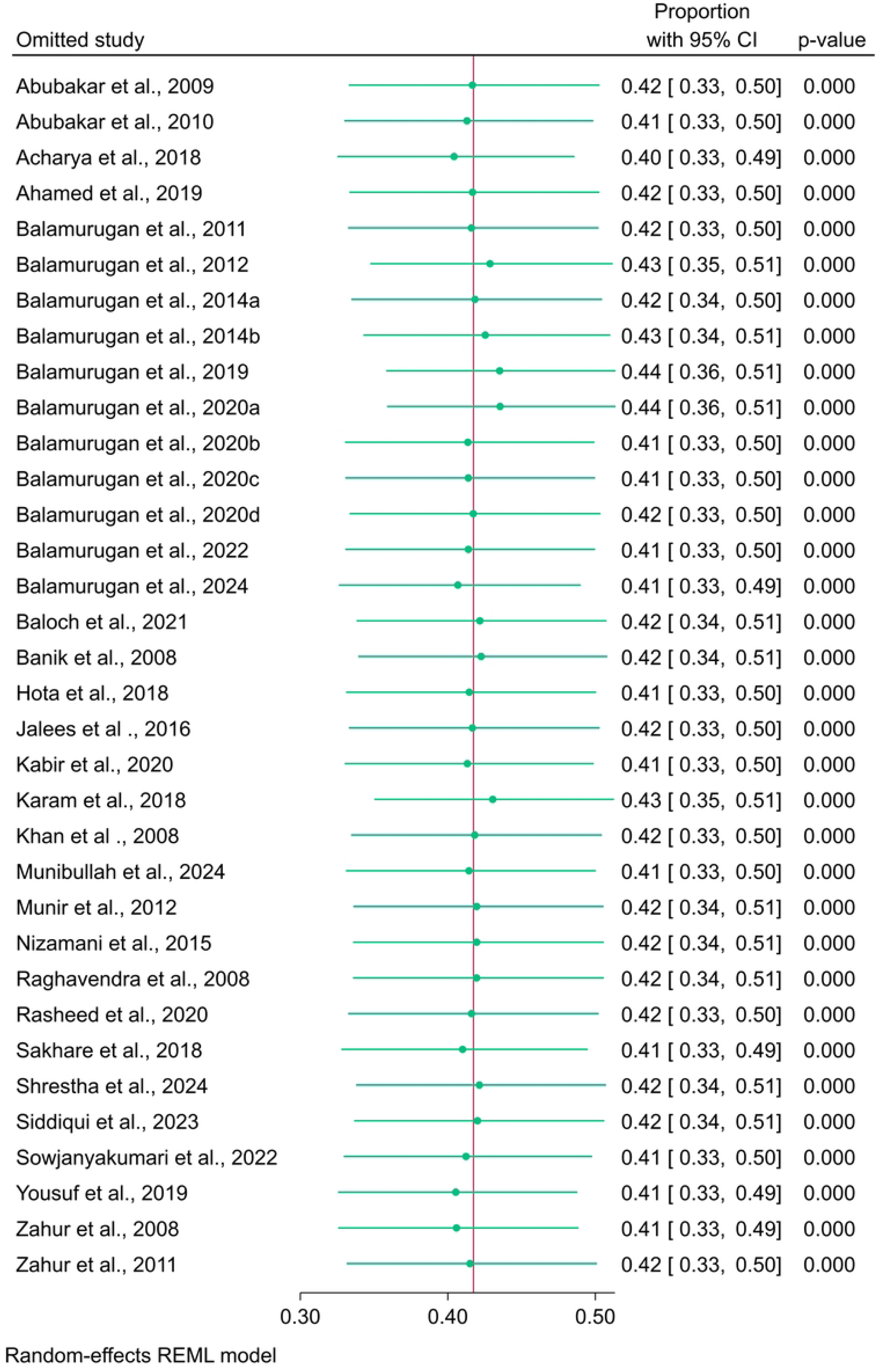
Leave-One-Out Sensitivity Analysis Plot for the PPR in Goats. Each point represents the pooled prevalence estimate when the corresponding study is omitted.

**Fig 18:**
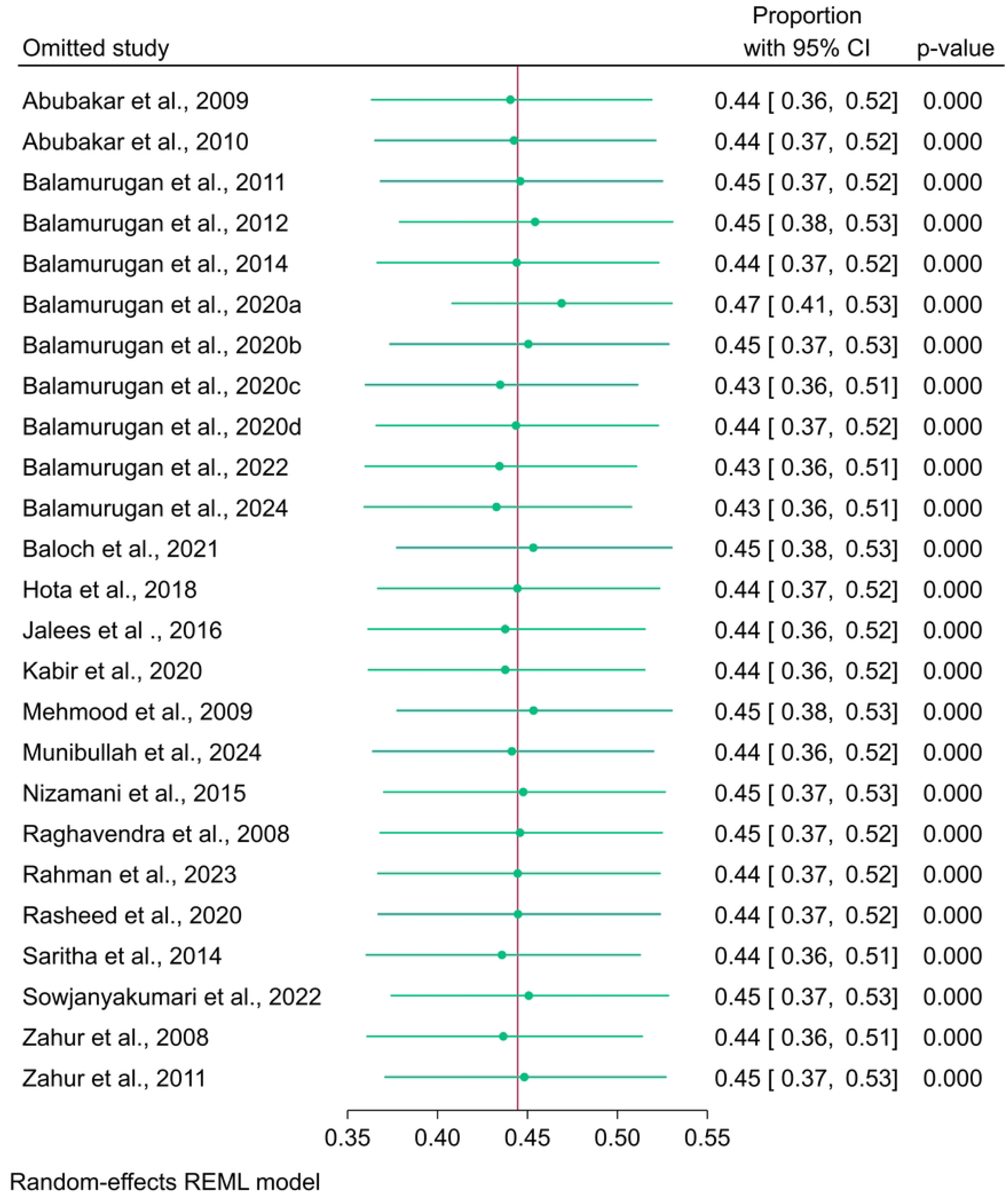
Leave-One-Out Sensitivity Analysis Plot for the PPR in Sheep. Each point represents the pooled prevalence estimate when the corresponding study is omitted.

**Table 4.** Univariate meta-regression of covariates.

| Covariates | *Sheep* |  |  |  | *Goat* |  |  |  | *Sheep and Goat* |  |  |  |
| --- | --- | --- | --- | --- | --- | --- | --- | --- | --- | --- | --- | --- |
| | $\beta$ | 95% CI | *p*-value | *R*<sup>2</sup> | $\beta$ | 95% CI | *p*-value | *R*<sup>2</sup> | $\beta$ | 95% CI | *p*-value | *R*<sup>2</sup> |
| Study year | 0.0031 | -0.0235 to<br>0.0297 | 0.819 | Nil | 0.0010 | -0.0306 to<br>0.0325 | 0.953 | Nil | 0.0017 | -0.0267 to<br>0.0301 | 0.905 | Nil |
| Country | 0.0406 | -0.2315 to<br>0.3127 | 0.770 | Nil | 0.0705 | -0.0843 to<br>0.2254 | 0.372 | Nil | 0.0473 | -0.0909 to<br>0.1856 | 0.502 | Nil |
| Vaccination | 0.4704 | 0.1411 to<br>0.7997 | **0.005** | 23.39 | 0.3297 | -0.1796 to<br>0.8390 | 0.205 | 1.82 | 0.3521 | -0.0764 to<br>0.7806 | 0.107 | 4.33 |
| Sample Size | 0.0001 | -0.0001 to<br>0.0003 | 0.456 | Nil | 2.48e<sup>-6</sup> | -0.0001 to<br>0.001 | 0.966 | Nil | 0.00001 | -0.0001 to<br>0.0001 | 0.757 | Nil |
| Positive<br>Cases | 0.0003 | -0.00002 to<br>0.0006 | 0.065 | 9.38 | 0.0002 | -0.0001 to<br>0.0004 | 0.216 | 1.57 | 0.0001 | -0.0001 to<br>0.0003 | 0.167 | 2.52 |

## 4. Discussion

*Peste des petits ruminants* (PPR) remains a major cause of mortality among millions of sheep and goats each year and poses a continuous threat to the livelihoods of small-scale farmers in South Asian countries. In South Asia, numerous cross-sectional studies and outbreak investigations have been conducted over the years. However, the reported sero-prevalence of PPR varies due to multiple influencing factors, such as study regions, sample size, vaccination status, time of surveys, and host animal species. Meta-analysis provides a useful approach to assess the impact of these variables by synthesizing data from diverse studies that differ in design, ecological context, and geographical setting. A well-conducted meta-analysis offers insights into disease dynamics that individual studies may fail to capture, thereby supporting more effective disease control strategies [71]. To our knowledge, this is the first quantitative meta-analysis focused on the seroprevalence of PPR in sheep and goats in South Asia. The analysis incorporates 37 (n=37) series of seroprevalence data collected between 2000 and 2025.

This study revealed a pooled seroprevalence of approximately 42.38% for PPR in sheep and goats across South Asia. According to the previous meta-analysis, the pooled seroprevalence of PPR in small ruminants was 39.2% [5], 40.1% [72] globally, and 27.7% in Ethiopia [9]. This variability can be attributed to several factors, including the geographical distribution of PPRV, the diagnostic techniques employed, the type and sources of the samples, the sampling strategies, the infection stages, the length of study periods, the host species of animals involved, and the number of samples analyzed. Differences in seroprevalence may also be influenced by regional ecological characteristics, such as climatic conditions, patterns of human settlement, hygiene standards, and socioeconomic practices [9].

The estimated overall seroprevalence revealed significant variation between species, with sheep showing a higher rate than goats. These findings align with many previous studies reported elsewhere [18,73–75]. In contrast, few studies have shown that goats have a higher seroprevalence than sheep [76–78]. Moreover, the findings of this study revealed that goats in Nepal, followed by those in Pakistan, presented greater seropositivity, whereas sheep in Pakistan presented a greater seropositivity rate than did those in other regions. Despite the biological differences between sheep and goats, a greater seroprevalence in one species than in the other may be influenced by factors such as the sampling strategy, population density or geographical distribution, husbandry practices, circulating virus strains, and/or other related variables [9]. PPRV may also exhibit a preference for infecting either sheep or goats based on environmental conditions, and the severity of the disease can vary between the two host species [79]. The subtotal pooled analysis indicated that, compared with Pakistan, Nepal had a greater seroprevalence of PPR compared to Pakistan in small ruminants. This might be the lack of published papers, especially in sheep. In Nepal, PPR outbreaks are often linked to the introduction of newly acquired animals from neighboring countries or foreign sources [80]. On the other hand, the high seroprevalence of PPR in sheep and goats in Pakistan and Bangladesh can be attributed to multiple factors. These factors include the cross-border movement of infected animals without adequate quarantine protocols, hot and humid climate conditions that favor the spread of PPRV, insufficient vaccination coverage and surveillance data, low awareness of PPR among smallholder farmers, and limited financial support for disease control initiatives in developing or underdeveloped countries [72]. India has adopted the PPR-Control Programme (PPR-CP) featuring a comprehensive ‘mass vaccination’ approach, which has significantly curtailed the number of outbreaks [81]. Despite being a neighboring country of India, Bangladesh continues to report a higher seroprevalence of PPR than India. Although a nationwide vaccination campaign was initiated in 2021, as recommended by WOAH, it aims to eradicate the disease by 2026 [82]. Consequently, the pooled estimated seroprevalence of PPR in these countries is comparatively low, reflecting the success of these control measures.

In this study, the overall pooled estimated sero-prevalence in sheep and goats was 57.5% in vaccination flocks in South Asia regions, suggesting a comparatively low level of herd immunity. Post-vaccination sero-prevalence rates were reported at 61.1% in Ethiopia [78] and 55.3% in Uganda [83], which supported the results demonstrated in the current study. The main reason for the low level of seroprevalence is a lack of vaccination coverage. PPRV antibodies detected in host animals may have been produced either through earlier vaccination or previous infection, as these antibodies are known to remain in the system for as long as 12 months [84]. Numerous studies included in this meta-analysis underscore the success of vaccination programs in reducing PPR outbreaks. Nevertheless, it is crucial to expand and standardize immunization efforts, particularly in regions highly susceptible to the disease. The provision of high-quality vaccines and the reinforcement of veterinary healthcare systems are fundamental to implementing effective PPR control strategies [85]. Furthermore, species-specific vaccination sero-monitoring based on pooled estimation indicated that sheep exhibited higher seroprevalence levels than goats in South Asia. This might be associated with many factors such as targeted vaccination coverage, species-specific immune response, effective husbandry practices, and population density. Prevaccination seroprevalence tends to vary across studies; however, postvaccination data consistently show that goats often exhibit antibody levels equal to or higher than those observed in sheep [86,87]. Although there is no definite cause of these variations, factors such as age, sex, and geographic region can affect seroprevalence rates. However, the difference between host species remains consistent across studies [26]. To improve immunity in both sheep and goats, an effective mass vaccination campaign is crucial, and the government should regularly assess the postvaccination seromonitoring.

### 4.1 Implications

The findings of this review and meta-analysis have significant implications for the eradication and management of PPR in South Asian countries recommended by the WOAH. The reliance on c-ELISA has led to an underestimation of the actual seroprevalence in these regions, because it has limited sensitivity. Additionally, the lack of vital information on host factors such as age, sex, and breed restricts the identification of risk groups and tailoring targeted interventions. The absence of published data further limits the understanding of the disease epidemiology. These findings demonstrate the urgent need for vast, recent, well-designed, and large-scale cross-sectional studies as well as the integration of novel techniques to improve disease monitoring and detection. Advanced data quality and assimilation of unpublished data into future meta-analyses and reviews will enhance effective and evidence-based policies for the eradication and prevention of PPR in South Asia and similar endemic regions in the world.

### 4.2 Limitations

This study has several limitations. Many of the studies lacked comprehensive data on key variables such as breed, age, and sex of the animals. The exclusion of unpublished data from the meta-analysis further restricts its ability to accurately represent the true seroepidemiological situation of PPR in the region. Furthermore, many of the included reports were outbreak-based, and due to the limited availability of cross-sectional studies, the analysis spans a lengthy timeframe from 2000 to 2025. As a result, the findings may not fully capture the current situation of the disease, the PPR in South Asia.

## 5. Conclusion

This study revealed that the pooled seroprevalence estimation of the disease, the PPR, is high in Nepal and low in India, even though a greater degree of variability was observed among the studies, between regions, and associated risk factors. The disease was highly prevalent in vaccinated sheep and goats, indicating that vaccination status has a major effect on seroprevalence. These findings suggest that screening tests for PPR and effective preventive and eradication measures should be routinely carried out in sheep and goat flocks in regions with high disease prevalence to control outbreaks and improve animal productivity. This meta-analysis highlights the need to enhance monitoring surveillance, improve vaccination coverage, and address the risk factors linked to disease transmission. To effectively implement control and eradication measures and reduce the burden of PPR, collaboration among government agencies, veterinary authorities, and foreign organizations is needed. The results of the present study suggest that further studies should prioritize specific regions, improve the quality of data, and evaluate the effects of vaccination efforts on the incidence of PPR. However, epidemiological surveillance is needed to estimate disease status and eliminate this disease. Moreover, factors that contribute to heterogeneity in prevalence estimates should be handled appropriately in survey studies to accurately estimate the true extent of PPR

## Abbreviation

CI: Confidence Interval
ELISA: Enzyme-Linked Immunosorbent Assay
HI: hemagglutination inhibition
JBI: Joanna Briggs Institute
OIE: Office International des Epizooties
PPR: Pesti des Petits Ruminants
PPRV: Pesti des Petits Ruminants Virus
PRISMA: Preferred Reporting Items for Systematic Reviews and Meta-Analyses
WOAH: World Organization for Animal Health

## Clinical trials

Not Applicable

## Availability of Data and Materials

The datasets used and/or analyzed during the current study are available in the supplementary materials.

## Author Contribution

**Md Jisan Ahmed:** Conceptualization (lead), investigation (lead), data curation (lead), methodology (lead), software (lead), validation (equal), formal analysis and interpretation of data (lead), data visualization (lead), writing—original draft (lead), writing—review and editing (supporting), and project administration (lead); **Md Imran Hossain:** Data extraction and curation (supporting), writing—original draft (supporting), and validation (equal); **Md Arifur Rahman:** Data extraction and data curation (supporting), data validation (supporting), & writing—original draft (supporting); **Ritu Chalise and Prajwal Bhandari**: Conceptualization (supporting), Data Extraction and Curation (lead), writing—review and editing (supporting), and validation (equal); **Tahmina Sikder, Amina Khatun, and Delower Hossain**: Writing—review and editing (lead), and validation (supporting). All the authors read the full manuscript and agreed to its publication.

## Ethical approval and consent to participate

Not Applicable

## Consent for publication

Not Applicable

## Acknowledgment

The author would like to acknowledge the technical support of the Association of Coding, Technology, and Genomics (ACTG), Sher-e-Bangla Agricultural University (SAU) members, especially Kazi Estieque Alam and Md Ismile Hossain Bhuiyan. The authors show appreciation towards Dr. Md. Mahabbat Ali for his support.

## Supplementary materials

S1 Table 1. Quality Assessment of Selected Articles

